# Piezo2 tension sensitivity and its modulation by alternative splicing

**DOI:** 10.64898/2026.02.16.706133

**Authors:** Michael Sindoni, William Sharp, Jörg Grandl

## Abstract

Piezo2 is a force-gated ion channel that functions as a sensor of mechanical touch, proprioception, lung inflation, and gut transit. Human Piezo2 contains seven domains that are alternatively spliced in a tissue-specific fashion resulting in the expression of at least 22 distinct variants. Despite the relevance of Piezo2 in human physiology, its sensitivity to membrane tension, and how this fundamental biophysical property is affected by alternative splicing, are unknown. Here, we use cell-attached pressure-clamp electrophysiology combined with differential interference contrast microscopy to quantify the response of Piezo2 to membrane tension and identify the alternatively spliced exon 35 as a domain sufficient to confer high sensitivity to membrane tension and cellular indentation. We further show that physiological variants of Piezo2 sense mechanical forces with distinct sensitivities and dynamic ranges. Together, our findings rationalize how Piezo2 variants may fulfill distinct physiological functions required for somatosensation and interoception.

## Introduction

The ability of cells to rapidly detect mechanical stimuli is enabled by force-gated ion channels. In mammals, the excitatory force-gated channels Piezo1 and Piezo2 are required for numerous cellular and physiological processes^1–4^. With few exceptions, Piezo1 and Piezo2 are expressed in non-overlapping tissues and cell-types, where they fulfill distinct physiological functions^5^. Piezo1 is broadly expressed in many cell types throughout the body and is required for basic cell functions such as mitosis, differentiation, and volume regulation, as well as organ homeostasis including bone formation, angiogenesis, and wound healing^6–12^. On the contrary, Piezo2 expression is largely limited to sensory tissue such as neurons of the dorsal root ganglia (DRG), the trigeminus and vagus nerves, and Merkel cells in the skin, and required for somatosensation and interoception including the sense of light touch, proprioception, bladder stretch, lung inflation, and gut transit^13–20^. Since Piezo channels have no known endogenous agonist, their function is thought to be specific to force sensing. Taken together with their physiological specialization, this suggests that Piezo1 and Piezo2 may differ in their ability to detect and transduce mechanical forces. Indeed, measurements using cell-attached pressure stimulation (stretch) and cellular compression with an atomic force microscopy (AFM) cantilever revealed that Piezo1 and Piezo2 respond to a different degree in both assays^21,22^. However, how Piezo1 and Piezo2 compare in what is arguably the most important functional aspect for many force-gated ion channels, the sensing of membrane tension, has yet to be determined.

Piezo1 is known to be directly activated by lateral membrane tension with a half-maximal activation (*T_50_*) of ∼1.4 mN/m, making it the most sensitive of all known force-gated channels^23–25^. For comparison, the inhibitory force-gated ion-channels TREK-1 (*T_50_* = 6.4±0.2 mN/m), TREK-2 (*T_50_* = 5.8±0.1 mN/m), and TRAAK (*T_50_* = 4.4±0.2 mN/m), as well as the excitatory TMEM63A (*T_50_* = 5.5±0.1 mN/m) all require much higher membrane tension to open^26,27^. Even more so, the bacterial channel MscL is activated near the lytic tension of the bilayer (*T_50_* = 11.8±0.8 mN/m)^28,29^. Quantification of the tension-response relationship of an ion channel allows calculation of the Gibbs free energy (ΔG) of channel gating, and the area expansion (ΔA) upon activation. For Piezo1, this knowledge has informed the interpretation of structural data, enabled comparisons between studies using AFM, minimal fluorescence photon fluxes (MINFLUX) super-resolution imaging, and channel reconstitution in small unilamellar vesicles, and allowed evaluating a theoretical framework of Piezo tension sensing^30–35^. In contrast, the ability of Piezo2 to sense membrane tension has not yet been determined. It is even unclear if Piezo2 is gated by membrane tension (force-from-lipids), force transmitted through other proteins (force-from-tether), or both. Disruption of the cytoskeleton of cells expressing Piezo2 impairs indentation-induced (poke) currents while promoting pressure-induced (stretch) responses, suggesting that Piezo2 can be directly gated by membrane tension, but that it is also sensitive to force transmission via linkage to the cytoskeleton^36^.

A precise measurement of Piezo2 tension sensing has remained elusive because, upon cell-attached pressure stimulation, few patches yield currents that are sufficient for generating tension response curves^21,36–39^. However, a recently generated expression vector of codon-optimized human Piezo2 that contains a WPRE regulatory element enhances protein production; this construct yields increased stretch-induced currents, making tension measurements possible^40,41^. Another complicating aspect is that Piezo2 does not exist as a singular isoform but is subject to alternative splicing. Specifically, sequencing of human tissue by Szczot and colleagues revealed that seven exons are alternatively spliced, resulting in at least 22 different Piezo2 variants in human DRG neurons and lung tissue alone^42^. Importantly, the abundance of different Piezo2 variants is tissue specific: For example, splice ‘variant 2’ constitutes ∼50% of all Piezo2 transcripts in human lung tissue but accounts for only ∼2% of all Piezo2 transcripts in DRG neurons. Further, splice variants that contain exon 35 are preferentially expressed in low threshold mechanoreceptors (LTMRs) that are involved in sensing discriminatory touch compared to high-threshold neurons that sense mechanical pain. Of course, such distinct tissue specificity implies a physiologically relevant difference in channel function. Indeed, Szczot and colleagues demonstrated that exon 33 modulates mouse Piezo2 permeability and calcium sensitivity, and that exon 35 accelerates inactivation kinetics^42^. However, a comprehensive investigation of how splicing affects the sensitivity of Piezo2 to mechanical forces has not been performed. Here, we precisely measure the sensitivity of Piezo2 to membrane tension and systematically investigate how this important function is modulated by alternative splicing.

## Results

### Alternative splicing modulates Piezo2 tension sensitivity

To determine if alternative splicing has any role in regulating the tension sensitivity of human Piezo2, we generated two constructs: one construct that lacks all alternatively spliced exons, hPiezo2_min_, and one that contains all alternatively spliced exons, hPiezo2_max_ (**Figure 1A, Supplemental Figure 1**). Since we were skeptical that channels lacking exon 22, which contains transmembrane domains 18 and 19, would remain functional, we decided to retain this exon in both constructs. Indeed, separate testing confirmed that removal of exon 22 renders both hPiezo2_min_ and hPiezo2_max_ non-functional (**Supplemental Figure 2**). To determine the tension sensitivity of both constructs, we heterologously expressed them in Neuro2A-Piezo1ko cells, which were CRISPR-engineered to lack Piezo1 and demonstrated to be devoid of mechanically-activated currents^43^. Next, we performed cell-attached pressure-clamp electrophysiology while simultaneously imaging the membrane dome with differential interference contrast (DIC) microscopy (**Figure 1B**). Both constructs exhibited prominent mechanotransduction currents that were absent in our YFP control (**Figure 1C**). Although relatively few patches from both constructs had currents that were large enough for our analysis, we overcame this limitation by patching many cells and focusing our analysis on measurements with peak currents of at least 20 pA (∼45% and ∼41% of patches from cells expressing hPiezo2_min_ and hPiezo2_max_, respectively) (**Figure 1D**). This allowed us to generate tension-response curves for each individual patch that were fit with a Boltzmann function to determine the tension of half-maximal activation (*T_50_*) and slope (*k*) (**Figure 1C**). Mean tension-response relationships from all patches showed that hPiezo2_max_ responds to lower tension values as compared to hPiezo2_min_ (**Figure 1E**). In fact, the tension-response relationships of individual patches revealed that values for *T_50_* and *k* differ ∼2-fold (hPiezo2_min_: *T_50_* = 3.7±0.2 mN/m, *k* = 1.2±0.1 m/mN, n = 17; hPiezo2_max_: *T_50_* = 2.0±0.1 mN/m, *k* = 2.3±0.3 m/mN, n = 24) (**Figure 1F**). This experiment confirmed our hypothesis that alternative splicing modulates Piezo2 tension sensitivity.

**Figure 1.**
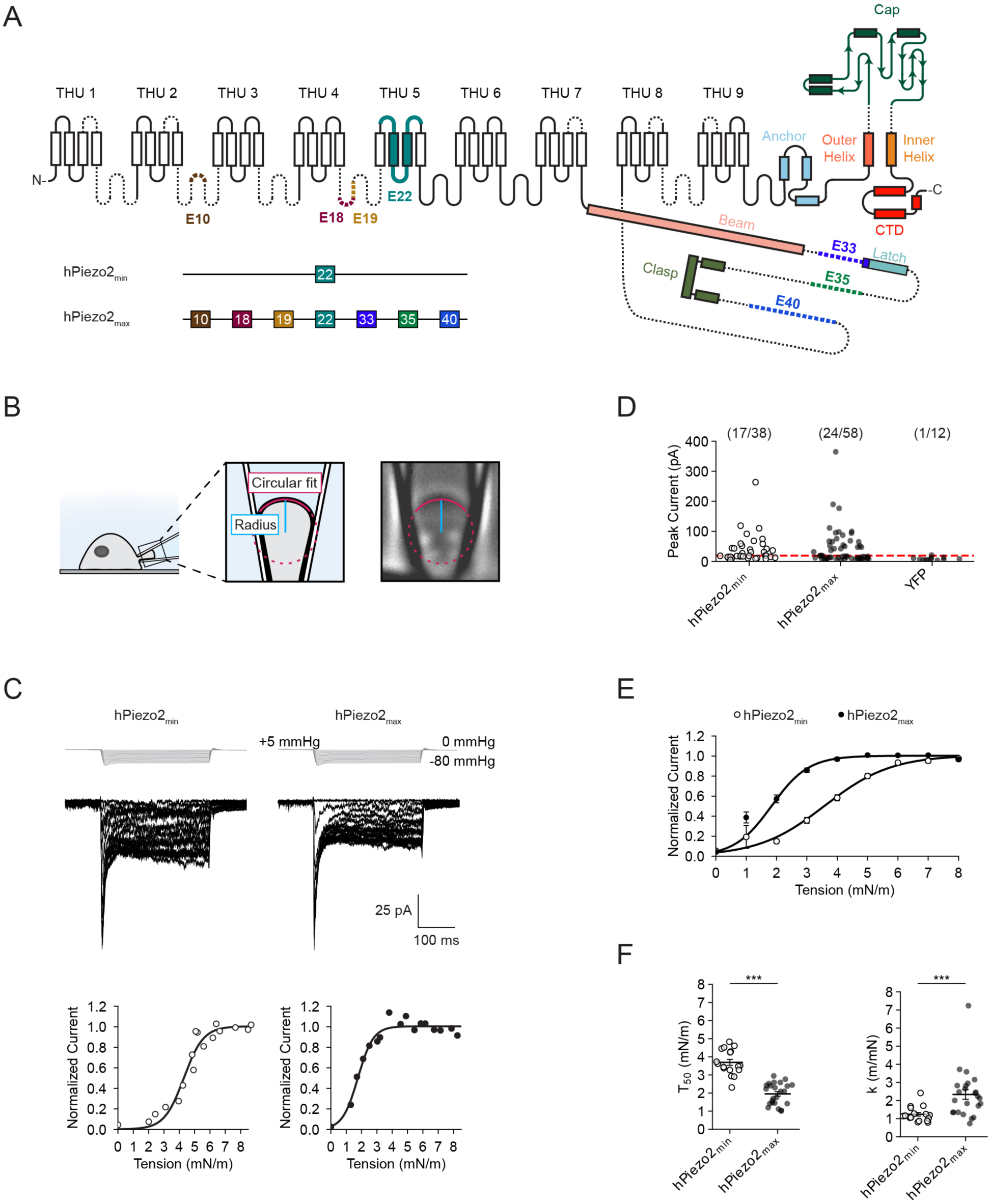
Tension responses of hPiezo2_min_ and hPiezo2_max_. ***A**, Top Schematic of human Piezo2 with key structural domains determined by the structure of mouse Piezo2. Dashed lines represent regions of the protein not resolved in mouse Piezo2. Alternatively spliced exons present in humans Piezo2 are colored (brown: exon 10, maroon: exon 18, yellow: exon 19, teal: exon 22, purple: exon 33, green: exon 35, blue: exon 40). Bottom: Schematic showing the alternatively spliced exons present in hPiezo2_min_ and hPiezo2_max_. **B**, Left: Schematic of combined cell-attached pressure-clamp electrophysiology and differential interference contrast (DIC) microscopy. Right: DIC microscopy image of a membrane dome inside a patch pipette at 400x magnification. A circular fit is applied to the membrane dome (red), which yields the radius of curvature (blue). **C**, Top: Pressure-step protocol from 0 to -80 mmHg (Δ = -5 mmHg) and representative currents recorded from Neuro2A-Piezo1ko cells heterologously expressing hPiezo2_min_ or hPiezo2_max_. Bottom: Normalized tension responses from the above recordings and Boltzmann fits. Current values are normalized to the plateau of the tension-response curve prior to normalization. **D**, Peak current amplitudes obtained with the above protocol from cells expressing hPiezo2_min_, hPiezo2_max_, or YFP alone. The red dashed line illustrates the 20 pA threshold above which patches were used for analyzing tension responses. The number of patches with peak currents > 20 pA and the total number of patches are shown above. **E**, Mean tension response curves for hPiezo2_min_ (n = 17) and hPiezo2_max_ (n = 24). Data are represented as normalized averages ± S.E.M with Boltzmann fits. **F**, Values for half-maximal activation (T_50_) and slope (k) from all individual patches shown in (E). Error bars indicate mean±S.E.M. Mean values for T_50_ and k are as follows: hPiezo2_min_ T_50_ = 3.7±0.2 mN/m, k = 1.2±0.1 m/mN; hPiezo2_max_ T_50_ = 2.0±0.1 mN/m, k = 2.3±0.3 m/mN. Significance was determined using Welch’s unpaired t-test (T_50_: t = 8.4, p < 0.0005; k: t = 3.8, p < 0.0005)*.

### Exon 35 confers high sensitivity to membrane tension

Next, we investigated which exons determine the difference in tension sensitivity we observed between hPiezo2_min_ and hPiezo2_max_. To do this, we pursued a systematic structure-function approach: we engineered six constructs in which we individually removed each exon from hPiezo2_max_ and six constructs where we individually added each exon to hPiezo2_min_, and measured their tension-response relationships (**Figure 2A**). All constructs exhibited peak current distributions similar to hPiezo2_min_ and hPiezo2_max_ (**Supplemental Figure 3**). We found that removing individual exons from hPiezo2_max_ did not alter tension-response curves for most constructs, indicating that they are not major contributors to the high tension sensitivity of hPiezo2_max_ (**Figure 2B, C**). One notable exception was hPiezo2_max-35_, the construct lacking only exon 35, because it exhibited increased *T_50_* values (*T_50_* = 2.5±0.3 mN/m, n = 10) as compared to hPiezo2_max_ (*T_50_* = 2.0±0.1 mN/m, n = 24), although this difference did not reach statistical significance. However, exon 35 clearly stood out when we analyzed constructs where individual exons were added to hPiezo2_min_ (**Figure 2D**). Specifically, hPiezo2_min+35_ exhibited a ∼1.5-fold difference in both *T_50_* and *k* values (*T_50_* = 2.4±0.2 mN/m, *k* = 2.1±0.4 m/mN, n = 15) as compared to hPiezo2_min_ (*T_50_* = 3.7±0.2 mN/m, *k* = 1.2±0.1 m/mN, n = 17), which was statistically significant (**Figure 2E**). This finding was specific to exon 35 because all other constructs where individual exons were added to hPiezo2_min_ did not exhibit statistically significant differences in *T_50_* and *k* values. Collectively, these results indicate that exon 35 is sufficient to confer high tension sensitivity to human Piezo2, and that exons 10, 18, 19, 33, and 40 individually do not have major roles in this function.

**Figure 2.**
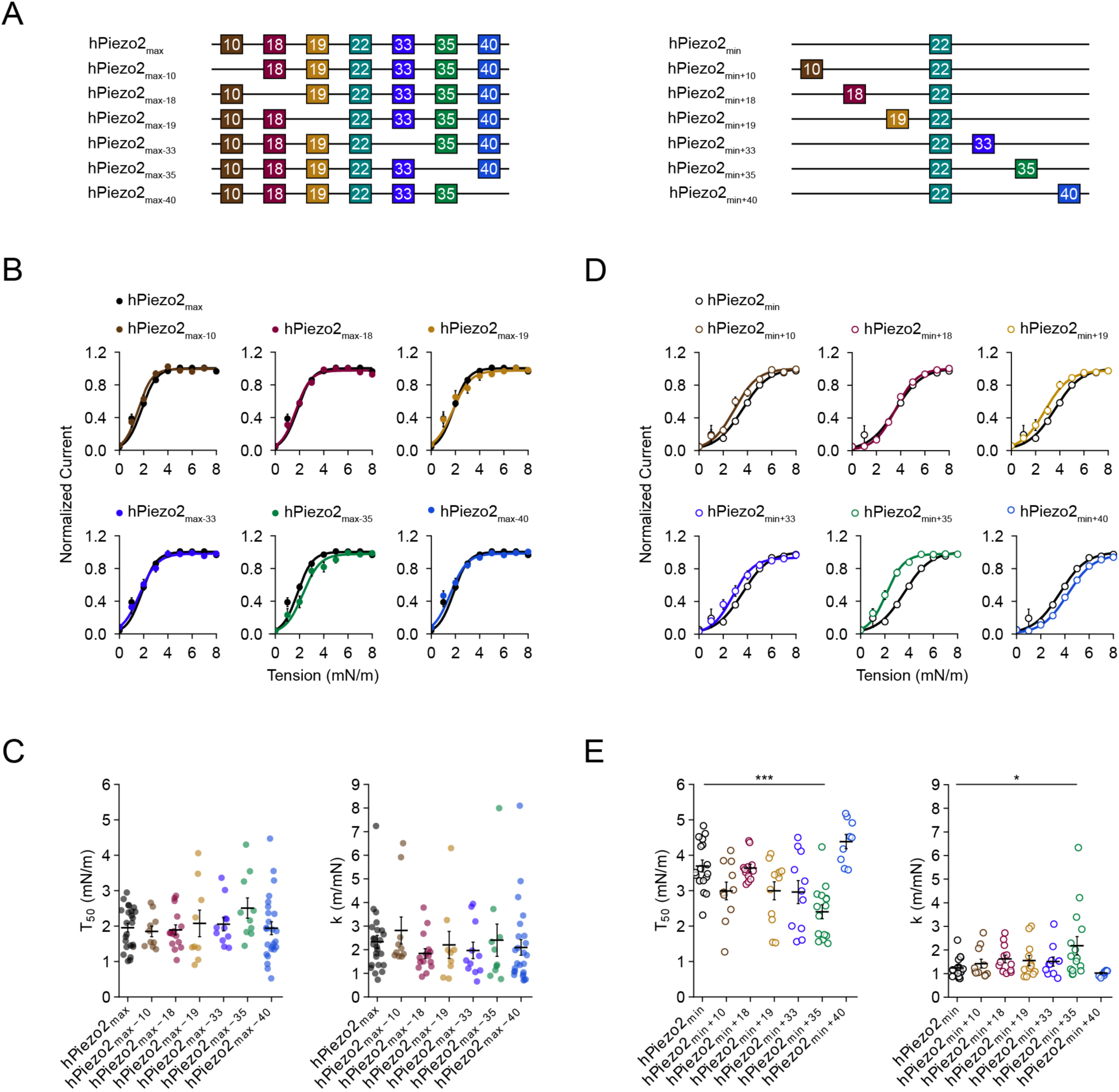
Tension responses of hPiezo2_min(+)_ and hPiezo2_max(-)_. ***A**, Schematic showing the alternatively spliced exons present and absent in hPiezo2_max(-)_ and hPiezo2_min(+)_ constructs. **B**, Mean tension response curves for hPiezo2_max_ (n = 24) and hPiezo2_max-10_ (n = 10), hPiezo2_max-18_ (n = 15), hPiezo2_max-19_ (n = 9), hPiezo2_max-33_ (n = 11), hPiezo2_max-35_ (n = 10), and hPiezo2_max-40_ (n = 24). Data are represented as normalized averages ± S.E.M with Boltzmann fits. **C**, T_50_ and k values of tension response curves from all individual patches shown in (B). Error bars indicate mean±S.E.M. Mean values for T_50_ and k are as follows: hPiezo2_max_ T_50_ = 2.0±0.1 mN/m, k = 2.3±0.3 m/mN; hPiezo2_max-10_ T_50_ = 1.9±0.2 mN/m, k = 2.8±0.6 m/mN; hPiezo2_max-18_ T_50_ = 1.9±0.2 mN/m, k = 1.8±0.2 m/mN; hPiezo2_max-19_ T_50_ = 2.1±0.4 mN/m, k = 2.2±0.6 m/mN; hPiezo2_max-33_ T_50_ = 2.1±0.2 mN/m, k = 2.0±0.4 m/mN; hPiezo2_max-35_ T_50_ = 2.5±0.3 mN/m, k = 2.4±0.7 m/mN; hPiezo2_max-40_ T_50_ = 1.9±0.2 mN/m, k = 2.1±0.3 m/mN. Significance was determined using a one-way ANOVA (F = 0.9, p = 0.48). **D**, Mean tension response curves for hPiezo2_min_ (n = 17) and hPiezo2_min+10_ (n = 11), hPiezo2_min+18_ (n = 12), hPiezo2_min+19_ (n = 12), hPiezo2_min+33_ (n = 11), hPiezo2_min+35_ (n =15), and hPiezo2_min+40_ (n =9). Data are represented as normalized averages ± S.E.M with Boltzmann fits. **E**, T_50_ and k values of tension response curves from all individual patches shown in (D). Error bars indicate average±S.E.M. Average values for T_50_ and k are as follows: hPiezo2_min_ T_50_ = 3.7±0.2 mN/m, k = 1.2±0.1 m/mN; hPiezo2_min+10_ T_50_ = 3.0±0.3 mN/m, k = 1.4±0.2 m/mN; hPiezo2_min+18_ T_50_ = 3.6±0.1 mN/m, k = 1.6±0.2 m/mN; hPiezo2_min+19_ T_50_ = 3.0±0.3 mN/m, k = 1.6±0.2 m/mN; hPiezo2_min+33_ T_50_ = 3.0±0.3 mN/m, k = 1.5±0.2 m/mN; hPiezo2_min+35_ T_50_ = 2.4±0.2 mN/m, k = 2.1±0.4 m/mN; hPiezo2_min+40_ T_50_ = 4.4±0.2 mN/m, k = 1.0±0.1 m/mN). Significance was determined using a one-way ANOVA (T_50_: F = 8.8, p < 0.0005; k: F = 2.6, p = 0.03) and Tukey’s HSD post-hoc comparison (hPiezo2_min_/hPiezo2_min+35_; T_50_, p < 0.0005; k, p = 0.03)*.

### Exon 35 confers low threshold to cell indentation

Piezo2 is also activated by cell indentation (poke), which has been reasoned to be a more relevant stimulus for its physiological functions^3,44^. We therefore wanted to know whether the effect of exon 35 is specific to pressure stimulation (stretch), or if it determines Piezo2 threshold to cell indentation. For this, we performed whole-cell indentation experiments on Neuro2A-Piezo1ko cells heterologously expressing hPiezo2_min_, hPiezo2_max_, hPiezo2_min+35_, or hPiezo2_max-35_, measured their indentation responses, and quantified indentation thresholds, the minimal indentation depth (*d*) after cell contact required to elicit mechanically activated currents (see Methods) (**Figure 3A, B, Supplemental Figure 4**). With this analysis we found that the indentation threshold was significantly higher for hPiezo2_min_ (*d* = 5.2±0.5 μm, n = 16) as compared to hPiezo2_max_ (*d* = 3.8±0.3 μm, n = 14), which readily confirms that splicing has a general role in modulating Piezo2 mechanical sensitivity (**Figure 3C**). Moreover, the indentation threshold of hPiezo2_min+35_ (*d* = 3.8±0.1 μm, n = 16) was significantly lower than that of hPiezo2_min_, but identical to that of hPiezo2_max_ (**Figure 3C**). Conversely, the indentation threshold of hPiezo2_max-35_ (*d* = 4.8±0.3 μm, n = 16) was increased compared to hPiezo2_max_, although this difference did not reach statistical significance (**Figure 3C**). Taken together, these results demonstrate that exon 35 is sufficient to confer not only high sensitivity to membrane tension, but also high sensitivity to cell-indentation.

**Figure 3.**
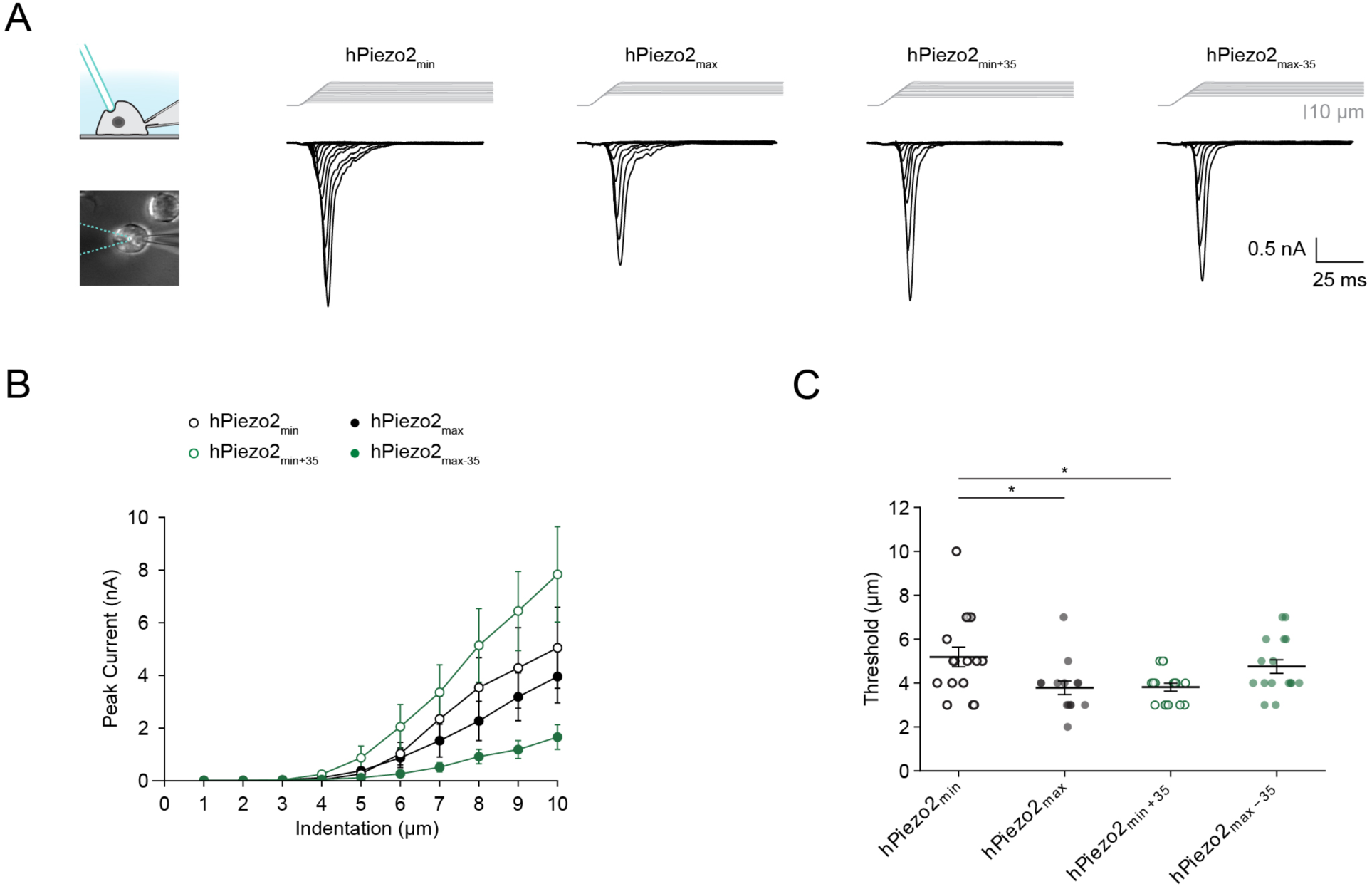
Indentation responses of hPiezo2_min_, hPiezo2_max_, hPiezo2_min+35_, and hPiezo2_max-35_. ***A**, Left: schematic of experimental setup and brightfield image showing mechanical poke stimulation of a cell using a blunt glass probe (blue) and recording in the whole-cell patch-clamp configuration. Right: Indentation-step protocol (Δ = 1 μm) and currents recorded from Neuro2A-Piezo1ko cells heterologously expressing hPiezo2_min_, hPiezo2_max_, hPiezo2_min+35_, or hPiezo2_max-35_. **B**, Average indentation response curves for hPiezo2_min_ (n = 16), hPiezo2_max_ (n = 14), hPiezo2_min+35_ (n = 16), and hPiezo2_max-35_ (n = 16). Data are plotted as mean ± S.E.M. **C**, Indentation threshold values for all individual patches shown in (B). Bars indicate the mean±S.E.M. Mean values for indentation threshold (d) are as follows: hPiezo2_min_ d = 5.2±0.5 μm; hPiezo2_max_ d = 3.8±0.3 μm; hPiezo2_min+35_ d = 3.8±0.1 μm; hPiezo2_max-35_ d = 4.8±0.3 μm. Significance was determined using a one-way ANOVA (F = 4.2, p = 0.009) and Tukey’s HSD post-hoc comparison (hPiezo2_min_/hPiezo2_max_, p = 0.03; hPiezo2_min_/hPiezo2_min+35_, p = 0.03).*

We also found that inactivation kinetics were nearly identical in hPiezo2_min_ (*τ = 4.2±0.7* ms, n = 16), hPiezo2_max_ (*τ = 4.0±0.9* ms, n = 14), hPiezo2_min+35_ (*τ = 3.7±0.4* ms, n = 16), and hPiezo2_max-35_ (*τ = 3.8±0.6* ms, n = 16) (**Supplemental Figure 5**). Thus, we conclude that alternative splicing of exon 35, or any other exon, does not affect the inactivation kinetics of human Piezo2.

### Physiological Piezo2 splice variants tile the tension range

Physiological membrane tension ranges from basal levels of 0.45 mN/m to near lytic tension ∼10 mN/m^45–47^. To what extent is this range dynamically transduced by Piezo1 and Piezo2 variants that exist physiologically? In mouse Piezo1, Geng and colleagues showed that removal of exon 30, which gives rise to its only known splice variant Piezo1.1, causes a 13.6 pS increase in unitary conductance and an 15.7 mmHg increase in pressure sensitivity^48^. We therefore analyzed single channel gating events and confirmed that both exons 30 and 33, which have modest sequence conservation (38% identity, 54% similarity), are necessary and sufficient to modulate the conductance of human Piezo1 and Piezo2, respectively (**Supplemental Figure 6**). Further, our own measurements confirmed that the modulation of pressure-sensitivity upon removal of exon 30 is conserved in human Piezo1.1 (**Supplemental Figure 7**), suggesting that it may also be more sensitive to membrane tension. Indeed, when measuring tension-response curves we found that human Piezo1.1 responds to lower tension values (*T_50_* = 1.7±0.1 mN/m, *k* = 3.0±0.4 m/mN, n = 8) as compared to human Piezo1 (*T_50_* = 2.4±0.1 mN/m*, k* = 2.5±0.3 m/mN, n = 21) (**Figure 4A-C, Supplemental Figure 8**). To visualize what these differences signify for transduction over the entire tension range, we calculated tension-tuning curves, the derivatives of tension-response curves (**Figure 4D**). The results show that human Piezo1 and Piezo1.1 both function in the lower range of physiological membrane tensions and that they transduce over a narrow dynamic range of ∼0-4 mN/m.

**Figure 4.**
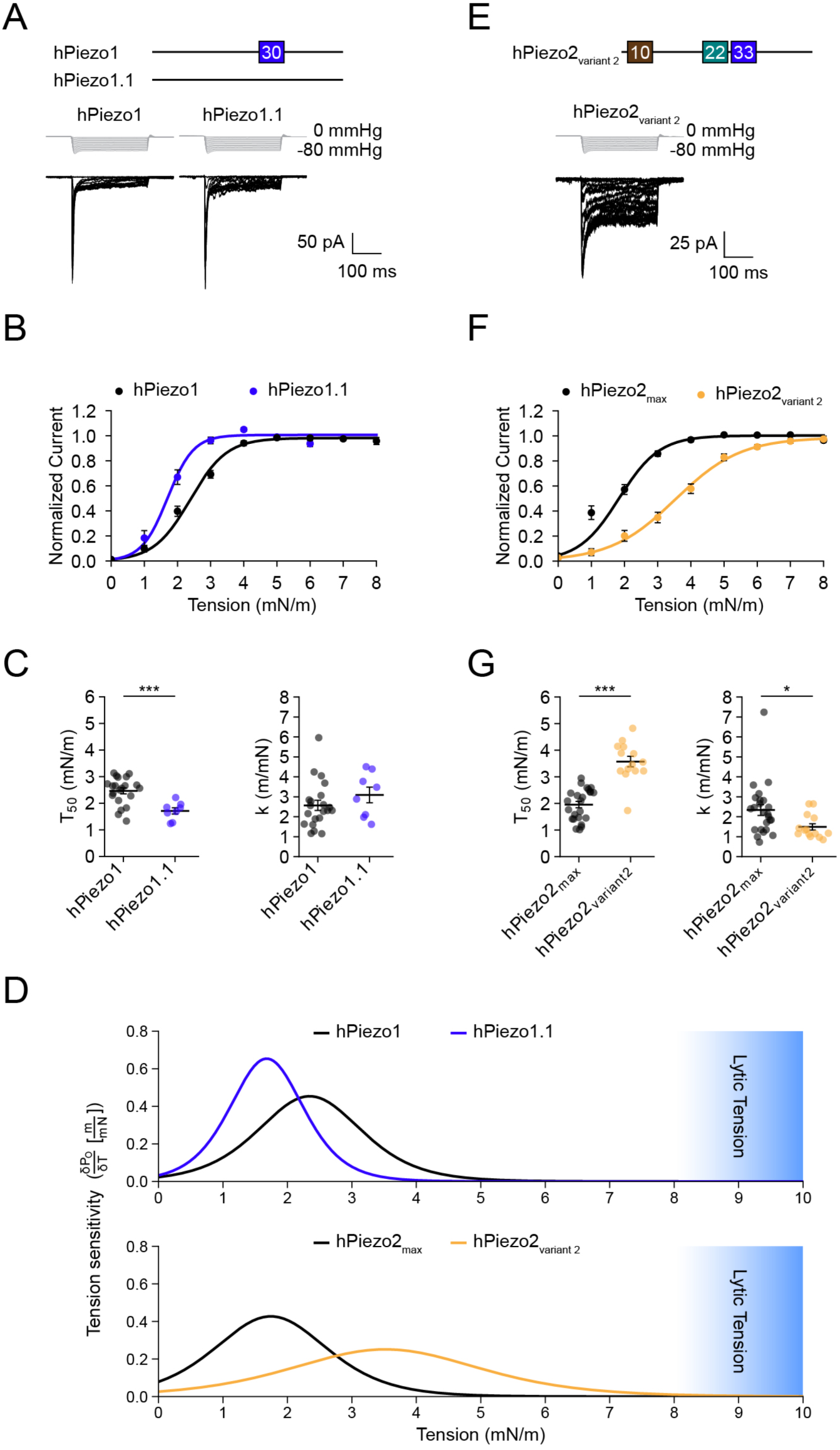
Comparison of physiological Piezo variant tension sensitivities. ***A**, Top: Schematic showing the alternatively spliced exons present in hPiezo1 and hPiezo1.1. Bottom: Pressure-step protocol from 0 to -80 mmHg (Δ = -5 mmHg) and representative currents recorded from Neuro2A-Piezo1ko cells heterologously expressing hPiezo1 or hPiezo1.1. **B**, Mean tension response curves for hPiezo1 (n = 21) and hPiezo1.1 (n = 8). Data are represented as normalized averages ± S.E.M with Boltzmann fits. **C**, T_50_ and k values from tension response curves from all individual patches from shown in (B). Error bars indicate mean±S.E.M. Mean values for T_50_ and k are as follows: hPiezo1 T_50_ = 2.4±0.1 mN/m, k = 2.5±0.3 m/mN; hPiezo1.1 T_50_ = 1.7±0.1 mN/m, k = 3.0±0.4 m/mN. Significance was determined using Welch’s unpaired t-test (T_50_: t = 4.7, p = 0.002; k: t = 1.1, p = 0.3). **D**, Tension sensitivity curves for hPiezo1 and hPiezo1.1 (top) and hPiezo2_min_, hPiezo2_max_, and hPiezo2 variant 2 (bottom). The range of tension that leads to membrane rupture (lytic tension) is indicated in blue. **E**, Top: Schematic showing the alternatively spliced exons present in hPiezo2 variant 2. Bottom: Pressure-step protocol from 0 to -80 mmHg (Δ = -5 mmHg) and representative currents recorded from Neuro2A-Piezo1ko cells heterologously expressing hPiezo2 variant 2. **F**, Mean tension response curves for hPiezo2_min_ (n = 17) and hPiezo2 variant 2 (n = 14). Data are represented as normalized averages ± S.E.M with Boltzmann fits. **G**, T_50_ and k values of tension response curves from all individual patches from shown in (F). Error bars indicate mean±S.E.M. Mean values for tension of half-maximal activation (T_50_) and slope (k) are as follows: hPiezo2_min_ T_50_ = 3.7±0.2 mN/m, k = 1.2±0.1 m/mN; hPiezo2_max_ T_50_ = 2.0±0.1 mN/m, k = 2.3±0.3 m/mN; hPiezo2 variant 2T_50_ = 3.6±0.2 mN/m, k = 1.5±0.2 m/mN. Significance was determined using a one-way ANOVA (T_50_: F = 43.4, p < 0.0005; k: F = 7.3, p < 0.0005) and Tukey’s HSD post-hoc comparison (T_50_: hPiezo2_max_/hPiezo2 variant 2, p < 0.0005; k: hPiezo2_max_ /hPiezo2 variant 2, p = 0.03).*

Regarding Piezo2, the most sensitive construct we identified in our systematic approach, hPiezo2_max_, is in fact an identified physiological variant that constitutes ∼16% of all human DRG transcripts^42^. On the contrary, the construct with the lowest sensitivity, hPiezo2_min_, has thus far not been detected in any tissue. We therefore focused instead on a variant that represents ∼50% of all transcripts in human lung tissue and was named ‘variant 2’ by Szczot and colleagues^42^. It contains only exons 10, 22, and 33, and importantly is lacking exon 35, suggesting that it may be tuned to sense high tension. In addition, variant 2 is noteworthy because it was the first human Piezo2 variant cloned, which makes it widely used across research labs^40,49,50^. We therefore measured the tension-response curve of variant 2 (*T_50_* = 3.6±0.2 mN/m, *k* = 1.5±0.2 m/mN, n = 14) and found that it differs substantially from hPiezo2_max_ (*T_50_* = 2.0±0.1 mN/m, *k* = 2.3±0.3 m/mN, n = 24), but is nearly identical to hPiezo2_min_ (*T_50_* = 3.7±0.2 mN/m, *k* = 1.2±0.1 m/mN, n = 17) (**Figure 4E-G, Supplemental Figure 8**). This finding agrees with the conclusion from our systematic structure-function approach that exons 10 and 33 alone do not confer high tension sensitivity. The corresponding tension-tuning curves show that hPiezo2_max_ transduces tension dynamically only between 0-4 mN/m, and the pronounced peak amplitude of its tuning curve highlights that it functions more like an on-off switch. In contrast, variant 2 transduces tension over a wide dynamic range of ∼1-7 mN/m, and the smaller peak of its tuning curve indicates that it does so much more gradually (**Figure 4D**). In other words, Piezo2_max_ is best suited to detect the lightest mechanical force, while variant 2 is optimal to encode force intensity. Overall, we conclude that alternative splicing of Piezo2 may tune tension sensing such that it tiles the range of physiological membrane tensions.

## Discussion

Splicing as a mechanism for modulating the sensitivity of sensory ion channels is a familiar concept. The heat sensor TRPV1 in vampire bats is alternatively spliced, altering its thermal threshold and sensitivity^51^. Similarly, the warm sensor TRPA1 is alternatively spliced in chemosensory neurons of fruit flies and mosquitoes to specifically reduce its thermosensitivity without affecting chemical sensing^52^. Our study adds to this knowledge by providing a rationale for how Piezo2 can fulfill a multitude of somatosensory and interoceptive functions, many of which may require distinct mechanical sensitivities and dynamic ranges.

Our systematic structure-function approach identified exon 35 as critical for conferring maximal tension sensitivity. This finding is remarkable, because sequencing and RT-PCR results from Szczot and colleagues show that Piezo2 transcripts that contain exon 35 are prominent in non-TRPV1 lineage neurons, which are a subset of low-threshold mechanoreceptors (LTMRs) known to be involved in sensing discriminative touch^42,53^. Similarly, Merkel cells in the skin are enriched in transcripts containing exon 35 and have been shown to mediate light touch sensing through Piezo2^13,14,19^. Conversely, transcripts containing exon 35 were nearly absent in TRPV1-lineage neurons which are involved in nociception^42,53^. Our study therefore provides one mechanism by which LTMRs and Merkel cells specialize to function as highly sensitive mechanoreceptors, whereas nociceptive neurons specialize in detecting mechanical forces with high threshold. Our results further show that variants lacking exon 35 have exceptionally broad dynamic ranges, making them well suited to transduce not only the presence of mechanical force, but also their intensities. Future long read sequencing studies on defined cell types could reveal which exact splice variants exist in the many tissues of the human body that rely on Piezo2 mechanotransduction.

The presence of exon 35 changes the mechanical sensitivity and dynamic range of Piezo2 substantially (∼1.5-fold). Interestingly, Geng and colleagues previously showed that differently spliced Piezo1 monomers can heterotrimerize into channels with intermediate conductances^48^. We therefore believe it should also be possible for Piezo2 monomers that contain exon 35 to assemble with monomers that lack it, thereby creating heteromeric channels with intermediate tension sensitivities and dynamic ranges. Further fine-tuning of Piezo2 mechanical sensing could be provided by known modulators of Piezo2. The auxiliary subunit MDFIC2 and the dietary fatty acid margaric acid both increase Piezo2-mediated mechanical indentation thresholds, while protein kinase A (PKA) reduces them^41,54–56^. Alternative splicing is distinct from these mechanisms in that it modulates the channel protein directly, thereby avoiding nonspecific effects on other signaling pathways. It will be interesting to test in the future if all Piezo2 splice variants are equally modulated by MDFIC2, margaric acid, and PKA, which may also provide insight into their still unknown mechanisms of action.

Our data also show that exons 10, 18, 19, and 40 do not play major roles in modulating tension sensing. Of course, it is possible that the contribution of each individual exon is too small to be detected by our assay, or that exons act synergistically to modulate this important function. Also, we found that these exons do not affect inactivation kinetics, and consistent with the original finding from Geng and colleagues, differences in conductance are conferred by exon 33 alone. We can therefore only speculate that exons 10, 18, 19, and 40 may provide other functions, such as sites for protein binding or post-translational modification, all of which warrants further investigation. Related to this, we do not know the logic underlying the splicing of exon 22, which we found is required for mechanically activated currents in both pressure-clamp and cell-indentation. Perhaps monomers lacking exon 22 may heterotrimerize with other spliced monomers to form functional channels, or act in a dominant-negative manner. Additionally, we found that exon 33 does not significantly affect the tension sensitivity of human Piezo2. This may seem striking because removal of the equivalent domain, exon 30, from human Piezo1, increases its tension sensitivity. However, the sequence conservation between both exons is modest (38% identity, 54% similarity), which may explain why exon 33 contributes less or not at all to modulating tension sensitivity. Alternatively, the limited resolution of our assay may also explain why we could have missed exon 33 as a domain important for tension sensing. Lastly, we did not observe that exon 35 alters Piezo2 inactivation kinetics, which differs from the findings of Szczot and colleagues^42^. This discrepancy may be a difference in species (human vs. mouse), cell lines (human HEK293T vs. mouse Neuro2A-Piezo1ko), or pipette buffers (nominal calcium vs. calcium free).

How might exon 35 affect the mechanical sensitivity of Piezo2? The tension-response curves we measured provide some insight: while the Gibb’s free energy (ΔG) of channel gating remains essentially unchanged between hPiezo2_min_ (4.6±0.4 k_B_T) and hPiezo2_min+35_ (4.6±0.5 k_B_T), the area expansion (ΔA) is nearly doubled between hPiezo2_min_ (5.1±0.4 nm^2^) and hPiezo2_min+35_ (9.0±1.6 nm^2^) (see Methods). Thus, our data predict that splice variants lacking exon 35 have a blade curvature that is less pronounced in the absence of tension, that membrane tension alone does not lead to complete blade flattening, or a combination of both. Structural studies, curvature measurements of small unilamellar vesicles containing single Piezo channels, and MINFLUX microscopy could all be used to test these predictions.

We believe this altered area expansion in channels lacking exon 35 could be caused by at least two distinct molecular mechanisms: First, a detailed theoretical framework of Piezo function by Haselwandter and MacKinnon showed that blade stiffness is a key contributing factor that determines the mechanical sensitivity of Piezos^35^. In fact, exon 35 could be well positioned to enhance overall blade stiffness, as it is part of a large intracellular loop that connects transmembrane helical units (THU) 7 and THU8 that are part of the blades. Unfortunately, cryo-electron microscopy (cryo-EM) studies have not yet resolved the structure of the intracellular loop that contains exon 35 and AlphaFold fails at predicting its structure with adequate confidence^57^. Second, exon 35 may provide a binding site for an accessory protein that modulates mechanical sensitivity. Recent work has shown that MDFIC2, which alters Piezo2 mechanical sensitivity, binds near the pore domain^41^. The intracellular loop that contains exon 35 is in principle close enough to interact physically with MDFIC2. Both hypotheses could be answered by additional structural data.

In conclusion, we measured Piezo2 tension sensitivity and demonstrated that it is modulated by alternative splicing. The identification of exon 35 as a regulatory domain for Piezo2 mechanical sensitivity will permit future studies to better understand its biophysical function and physiological roles.

## Materials and Methods

### Plasmid design

All human Piezo2 constructs were generated using the hPIEZO2-codon-optimized-pIRES2-mCherry-WPRE cDNA (a gift from Ardem Patapoutian^40^), also referred to as ‘variant 2’. Individual exons were added or removed using the Q5 mutagenesis kit (New England Bioscience) with non-overlapping primers (see Key Resources Table) according to the manufacture’s protocol. The human Piezo1.1 construct was generated using hPiezo1-pIRES2-EGFP cDNA (Ardem Patapoutian^58^) by removing exon 30 as previously described. All constructs were verified using overlapping primers to sequence the entire protein coding region (Genewiz).

### Cell culture

Neuro2A-Piezo1ko cells (a gift from Gary Lewin^43^) were cultured at 37°C and 5% CO_2_ in Minimum Essential Medium (Thermo Fisher Scientific) supplemented with 50 U/ml penicillin (Thermo Fisher Scientific), 50 mg/ml streptomycin (Life Technologies), 0.1 mM non-essential amino acids (Thermo Fisher Scientific), 1 mM Sodium Pyruvate (Thermo Fisher Scientific), and 10% fetal bovine serum (Clontech). For all overexpression experiments, cells were transiently transfected in six-well plates using Lipofectamine 2000 (Thermo Fisher Scientific) 40-48 hours before recording. For all human Piezo2 experiments, cells were transfected with 4 µg cDNA for individual human Piezo2 constructs and 1 µg cDNA for yellow fluorescent protein (YFP) or 4 µg cDNA for YFP alone. For human Piezo1 experiments, cells were transfected with 4 µg cDNA for either human Piezo1, human Piezo1.1, or YFP. 16–24 hours before recording cells were reseeded onto P50G-0-30-F glass-bottomed dishes (MatTek Corporation) coated with poly-L-lysine and laminin (Sigma) and cultured in the above-described media with 20 μM ruthenium red (Sigma) added.

### Electrophysiology

Cell-attached patch-clamp recordings were performed at room temperature using an EPC10 amplifier and Patchmaster software (HEKA Elektronik). Data were sampled at 10 kHz and filtered at 2.9 kHz. Borosilicate glass pipettes (1.5 OD, 0.86 ID; Sutter Instrument Company) had a resistance of 1.5–3 MΩ when filled with pipette buffer solution (in mM): 130 NaCl, 5 KCl, 10 HEPES, 10 TEACl, 1 CaCl_2_, 1 MgCl_2;_ pH = 7.4 with NaOH, osmolarity = 302 mOsm/L). The bath solution was (in mM): 140 KCl, 10 HEPES, 1 MgCl_2_, 10 glucose; pH = 7.3 with KOH, osmolarity = 286 mOsm/L. Pressure was controlled with an HSPC-2-SB high speed pressure clamp system (ALA Scientific). For tension-response measurements, pipettes were positioned at an angle of ∼15° with respect to the glass-bottomed dishes to optimize image contrast. Patches were held at -80 mV for the entire protocol. To mechanically stimulate the membrane, first, a positive pressure step of +5 mmHg was applied for five seconds to minimize basal membrane tension upon patch formation. Next, negative pressure steps from 0 to -80 mmHg (Δ = -5 mmHg) were applied for 300 ms. All pressure steps were separated by 10 s to allow for recovery from inactivation. A loss of gigaseal or membrane density in the images was interpreted as patch rupture and subsequent pressure steps were not analyzed. Single-channel conductance measurements were performed using the same buffers and equipment as described above. Pipettes were positioned at an angle of ∼45° with respect to the glass-bottomed dishes. Single-channel openings were recorded for 10 s using a voltage-step protocol from -60 mV to -100 mV (Δ = -20 mV) and -5 mmHg of negative pressure to elicit channel opening for 10 s. Voltage steps were separated by 10 s to allow for recovery from inactivation. All cell-attached electrophysiological recordings were only analyzed from patches with seal resistances >1 GΩ and leak currents <50 pA. N denotes the number of individual cells patched, with no cell being patched more than one time.

Whole-cell recordings were performed using the amplifier and program described above. Borosilicate glass pipettes (1.5 OD, 0.86 ID; Sutter Instrument Company) had a resistance of 1.5–2.5 MΩ when filled with pipette buffer solution (in mM): 133 CsCl, 2 MgCl_2_, 10 HEPES, 1 EGTA, 4 MgATP, 0.4 Na_3_GTP_;_ pH = 7.4 with CsOH, osmolarity = 282 mOsm/L). The bath solution was (in mM): 150 NaCl, 2 KCl, 1 MgCl_2_, 2.5 CaCl_2_, 10 glucose, 10 HEPES; pH = 7.4 with NaOH, osmolarity = 318 mOsm/L. Patch pipettes were positioned at an angle of ∼45° with respect to the glass-bottomed dishes and patches were held at -80 mV for the entire protocol. To apply mechanical stimulation, cells were indented with a fire-polished glass pipette (tip diameter ∼1-3 μm) using an amplifier-controlled piezo-electric driver and E-625 Piezo Servo Controller (Physik Instrumente) operating in closed-loop mode. The probe was positioned above the cell and advanced at 1 μm/ms at an 80° angle. Indentation depth was increased in 1 μm increments, and the probe was held at each indentation for 500 ms, before being retracted to its resting position at a speed of 1 μm/ms. All indentation steps were separated by 10 s to allow for recovery from inactivation. The indentation step protocol was continued until patch rupture. Electrophysiological recordings were only analyzed from patches with compensated series resistances <15 MΩ, leak currents <100 pA, and capacitance <40 pF. N denotes the number of individual cells patched, with no cell being patched more than one time.

### Image acquisition

For tension-response measurements, images of the membrane dome inside the patch pipette were acquired using an Eclipse Ti (Nikon) inverted microscope and Plan Apo (100x) differential interference contrast (DIC) oil objective (Nikon). Images were further magnified with a 4x relay lens (Nikon) and recorded with a CoolSNAP ES2 camera (Photometrics). During imaging, the focal plane was manually adjusted to maintain focus on the membrane dome. Images were acquired using either Nikon NIS-Elements (Melville, NY) or the ‘Micro-Manager’ (Windows 64-bit, version 2.0.0) open-source software^59^ at a rate of 10 Hz (100 ms exposure). Pixels were binned 2x2, resulting in a resolution of 0.0325 μm/pixel. For indentation-response measurements, cells were imaged using the same camera and microscope described above with an S Plan Fluor 40x objective (Nikon) and acquired at 10Hz using ‘Micro-Manager’ software.

### Quantification and statistical analysis Tension-response measurements

All electrophysiology data were analyzed with a custom script using Python version 3.8.5 (<u>github.com/GrandlLab</u>). Normalized peak currents were determined the following way. First the mean current measured before the onset of the pressure step was baseline subtracted from each recording. Second, the peak current was calculated as the mean of the absolute peak current and the two data points preceding and following it. Only patches with a maximal peak current >20 pA were included in tension-response analyses. Individual pressure-step peak currents that showed rundown (>30% reduction in subsequent peak currents) were excluded from further analyses. To determine normalized peak currents, a pressure–response curve was generated and fit with the following Boltzmann function:

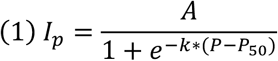

where *I_p_* is the peak current, *A* is the plateau amplitude, *k* is the slope, *P* is the pressure applied at each pressure step, and *P_50_* is the pressure of half-maximal activation. Each peak current was then divided by the plateau amplitude *A* of the fit.

Imaging data were analyzed with a custom script using Python version 3.8.5 (<u>github.com/GrandlLab</u>). First, at each pressure step, the image at which the membrane achieved the greatest degree of curvature was manually selected. Second, an ROI containing the membrane dome and its contact points with the pipette walls was manually selected. Next, the mean pixel intensity was calculated for segments of five pixels in each single column. Next, every column was re-analyzed to 1) manually discard pixel columns that are not part of the membrane dome, 2) autonomously identify the segment with the lowest mean pixel intensity, 3) identify 25 pixels neighboring the center pixel of this segment, and 4) fit the intensity values of the 25 pixels of each column with the following Gaussian function:

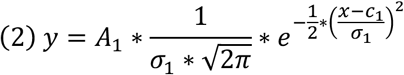

where x is the center location of each bin, y is the count for each bin, and *A_1_* is the amplitude, *C_1_* is the center value, and *σ_1_* is the standard deviation of each Gaussian fit. Values of peak location and standard deviation obtained from the Gaussian fits of each column were then used for a weighted circular fit to obtain membrane dome curvature radius (*r*) using the following equation:

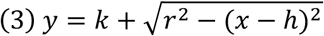

where *x* and *y* are the coordinates of the circle circumference, *h* and *k* are the coordinates for the circle center, and *r* is the radius. Lastly, segments of fitted circles were digitally appended to the original image to allow for visual inspection of the fit. Images with poor circular fits, defined by a relative standard deviation of the radius 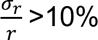, were excluded from the dataset, except for images where pressure was 0 mmHg. If more than two images were excluded, the entire patch was excluded from further analysis. Lastly, membrane tension was determined using Laplace’s equation:

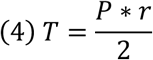

where *T* is membrane tension, *P* is the pressure applied through the patching pipette, and *r* is the radius of curvature calculated above.

Average tension-response curves were generated by normalizing peak currents, binning tension values in increments of 1 mN/m, and fitting them with Boltzmann functions:

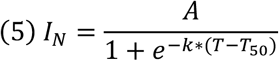

where *I_N_* is the normalized peak current, *A* is the plateau amplitude, and *T* is the tension elicited at each pressure step.

For each construct the mean±S.E.M. for *T_50_* and *k* were calculated and used for statistical analysis.

For individual patches, values for tension of half-maximal activation (*T_50_*) and slope (*k*), were generated by normalizing peak currents, without binning tension values, and fitting with Boltzmann functions (5).

### Indentation threshold measurements

For whole-cell poke experiments, indentation threshold values for individual patches were determined in the following way: First, videos were recorded during the experiment. The indentation depth at which the glass probe first contacted the cell was identified offline by inspecting video recordings, and all measurements were subsequently aligned to that indentation. Second, peak current at each indentation depth was determined as described above. Third, a histogram of peak currents for all indentation depths and patches was generated and we determined that a threshold of 10 pA was ideal to define electrical responsiveness to mechanical stimulation (**Supplemental Figure 4D**). Lastly, for each construct the indentation threshold (*d*) ± S.E.M. was calculated and used for statistical analysis.

### Inactivation kinetics (tau) measurements

To determine inactivation tau (*τ*), the current from the final indentation step of each patch was fit with the following single-exponential function:

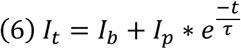

where *I_t_* is the is the current at a given time, *I_b_* is the baseline current, *I_p_* is the peak current elicited during the indentation step, and *t* is time. For each construct the tau (*τ*) ± S.E.M. was calculated and used for statistical analysis.

### Conductance measurements

For single-channel conductance experiments, the slope conductance for individual patches was determined the following way. First, 3-5 single-channel opening events were isolated for each voltage-step and current histograms were generated and fit with the following double Gaussian equation:

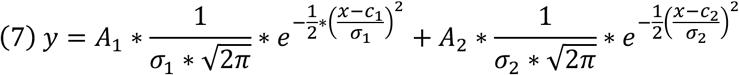

where x is the center location of each bin, y is the count for each bin, and *A_1/2_* are the fit amplitudes, *C_1/2_* are the center values, and *σ_1/2_* are the standard deviations of each Gaussian fit. Second, the difference between *C_1_* and *C_2_* was used to determine the unitary current *i_u_*. Third, a current-voltage plot was generated and the data fit with the following linear function:

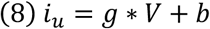

where *i_u_* is the unitary current, *g* is the slope conductance, *V* is voltage, and *b* is the y-intercept. For each construct the mean slope conductance (*g*) ± S.E.M. was calculated and used for statistical analysis.

### Pressure-response measurements

All electrophysiology data were processed and analyzed as described above. Normalized peak currents were determined as they were above using equation (1). To determine the pressure of half-maximal activation (*P_50_*) and slope (*k*), normalized pressure–response curves were generated and fit with the following Boltzmann function:

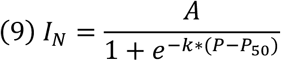

where *I_N_* is the normalized peak current, *A* is the plateau amplitude, and *P* is the pressure at each pressure step. For each construct the average±S.E.M. for *P_50_* and *k* were calculated and used for statistical analysis.

### Tension sensitivity curves

Average tension-response curves for all patches were generated for each construct as follows: Normalized peak currents were binned in increments of 1 mN/m. Histograms were fit with a weighted Boltzmann fit (equation 5) using the mean currents and their standard deviations to yield values for *T_50_*, *k* and *A*. Tension-sensitivity curves were generated by calculating the first derivative of the Boltzmann equation (equation 4):

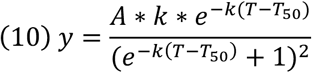

where *T* is tension, *A* is the amplitude, *T_50_* is the tension of half-maximal activation, and *k* is the slope of the tension-response Boltzmann fit.

### Thermodynamic parameters of channel gating

Values for Gibbs free energy (ΔG) and channel area expansion (ΔA) were determined using the following equations:

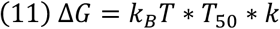

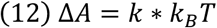

, where *k*_B_ is the Boltzmann constant and *T* is the absolute temperature (300 K).

### Statistical analyses

Specific statistical analysis and sample sizes for each experiment can be found in their respective figure legends. All summary data are represented as average±S.E.M. Statistical significance is represented the following way: * = p < 0.05; ** = p < 0.0005; *** = p < 0.0005. For all experiments, day matched controls were used. To prevent bias in identifying first contact, all indentation threshold experiments and analysis were performed double blinded.

## Author Contributions

M.S. and W.S. designed and performed experiments and analyzed data. M.S. and J.G. wrote the manuscript.

## Declaration of Interests

The authors declare no competing interests.

## Acknowledgments

We thank Dr. Amanda H. Lewis, Jialin Wu, and Dr. Marie E. Cronin for feedback on the manuscript. Funding was provided by NIH F31NS139449 and NIH 5R01NS110552.

## Key Resources Table

| REAGENT or RESOURCE | SOURCE | IDENTIFIER |
| --- | --- | --- |
| **Experimental models: Cell lines** |  |  |
| Neuro2A-Piezo1ko cells | Gift of Gary Lewin; Moroni et al.<sup>43</sup> | N/A |
| **Oligonucleotides** |  |  |
| See Table S1-5 for primers to generate hPiezo2 and hPiezo1 constructs | This paper | N/A |
| **Recombinant DNA** |  |  |
| Human PIEZO2-pIRES2-mCherry-WPRE (hPiezo2<sub>variant 2</sub>) | Villarino et al.<sup>40</sup> | Construct #1 |
| Human PIEZO2<sub>max</sub>-pIRES2-mCherry-WPRE (hPiezo2<sub>max</sub>) | This paper | Construct #8 |
| Human PIEZO2<sub>max-10</sub>-pIRES2-mCherry-WPRE (hPiezo2<sub>max-10</sub>) | This paper | Construct #12 |
| Human PIEZO2<sub>max-18</sub>-pIRES2-mCherry-WPRE (hPiezo2<sub>max-18</sub>) | This paper | Construct #13 |
| Human PIEZO2<sub>max-19</sub>-pIRES2-mCherry-WPRE (hPiezo2<sub>max-19</sub>) | This paper | Construct #14 |
| Human PIEZO2<sub>max-22</sub>-pIRES2-mCherry-WPRE (hPiezo2<sub>max-22</sub>) | This paper | Construct #15 |
| Human PIEZO2<sub>max-33</sub>-pIRES2-mCherry-WPRE (hPiezo2<sub>max-33</sub>) | This paper | Construct #16 |
| Human PIEZO2<sub>max-35</sub>-pIRES2-mCherry-WPRE (hPiezo2<sub>max-35</sub>) | This paper | Construct #17 |
| Human PIEZO2<sub>max-40</sub>-pIRES2-mCherry-WPRE (hPiezo2<sub>max-40</sub>) | This paper | Construct #5 |
| Human PIEZO2<sub>min</sub>-pIRES2-mCherry-WPRE (hPiezo2<sub>min</sub>) | This paper | Construct #10 |
| Human PIEZO2<sub>min-22</sub>-pIRES2-mCherry-WPRE (hPiezo2<sub>min-22</sub>) | This paper | Construct #11 |
| Human PIEZO2<sub>min+10</sub>-pIRES2-mCherry-WPRE (hPiezo2<sub>min+10</sub>) | This paper | Construct #9 |
| Human PIEZO2<sub>min+18</sub>-pIRES2-mCherry-WPRE (hPiezo2<sub>min+18</sub>) | This paper | Construct #18 |
| Human PIEZO2<sub>min+19</sub>-pIRES2-mCherry-WPRE (hPiezo2<sub>min+19</sub>) | This paper | Construct #19 |
| Human PIEZO2<sub>min+33</sub>-pIRES2-mCherry-WPRE (hPiezo2<sub>min+33</sub>) | This paper | Construct #25 |
| Human PIEZO2<sub>min+35</sub>-pIRES2-mCherry-WPRE (hPiezo2<sub>min+35</sub>) | This paper | Construct #21 |
| Human PIEZO2<sub>min+40</sub>-pIRES2-mCherry-WPRE (hPiezo2<sub>min+40</sub>) | This paper | Construct #24 |
| Human Piezo1-pIRES2-EGFP (hPiezo1) | Albuisson et al.<sup>58</sup> | Construct #26 |
| Human Piezo1.1-pIRES2-EGFP (hPiezo1.1) | This paper | Construct #27 |
| pCAG-Yellow Fluorescent Protein (YFP) | https://www.addgene.org/11180/ | RRID:Addgene_11180 |
| **Software and algorithms** |  |  |
| Nikon NIS-Elements | https://www.microscope.healthcare.nikon.com/products/software/nis-elements | RRID:SCR_002776 |
| Micro-Manager (Windows 64-bit, version 2.0.0) | Edelstein et al.<sup>59</sup> | RRID:SCR_000415 |
| Python (version 3.8.5) | https://www.python.org/ | RRID:SCR_008394 |
| Custom electrophysiology analysis Codes | This paper | https://github.com/GrandLab |
| Custom image analysis Codes | This paper | https://github.com/GrandLab |
| **Other** |  |  |
| High-Speed Pressure Clamp (HSPC-2-SB) | ALA Scientific | N/A |
| EPC10 amplifier | HEKA Elektronik | RRID:SCR_018399 |
| CoolSNAP ES2 camera | Photometrics | N/A |
| E-625 Piezo Servo Controller | Physik Instrumente | N/A |

## Supplemental information

**Supplemental Figure 1.**
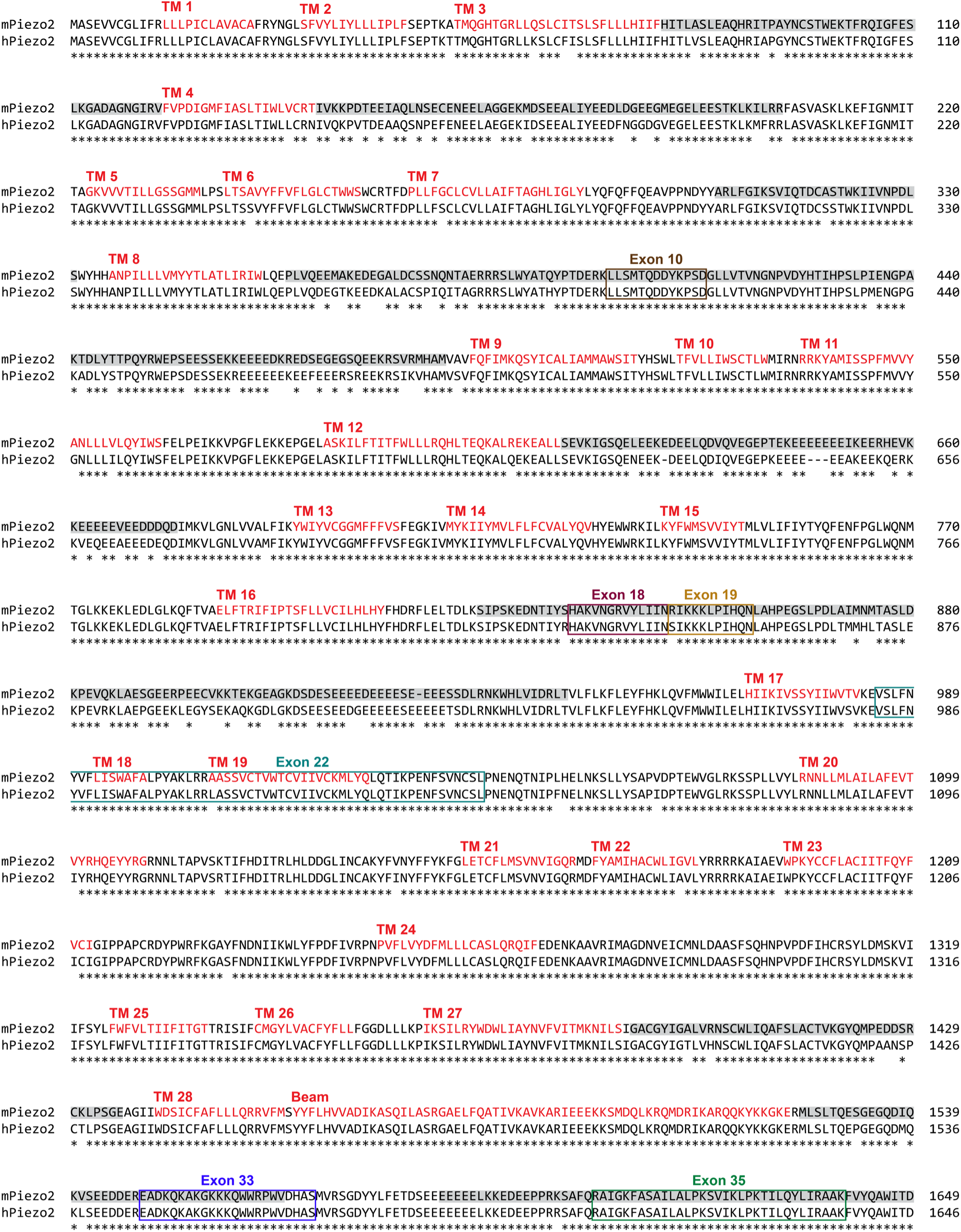
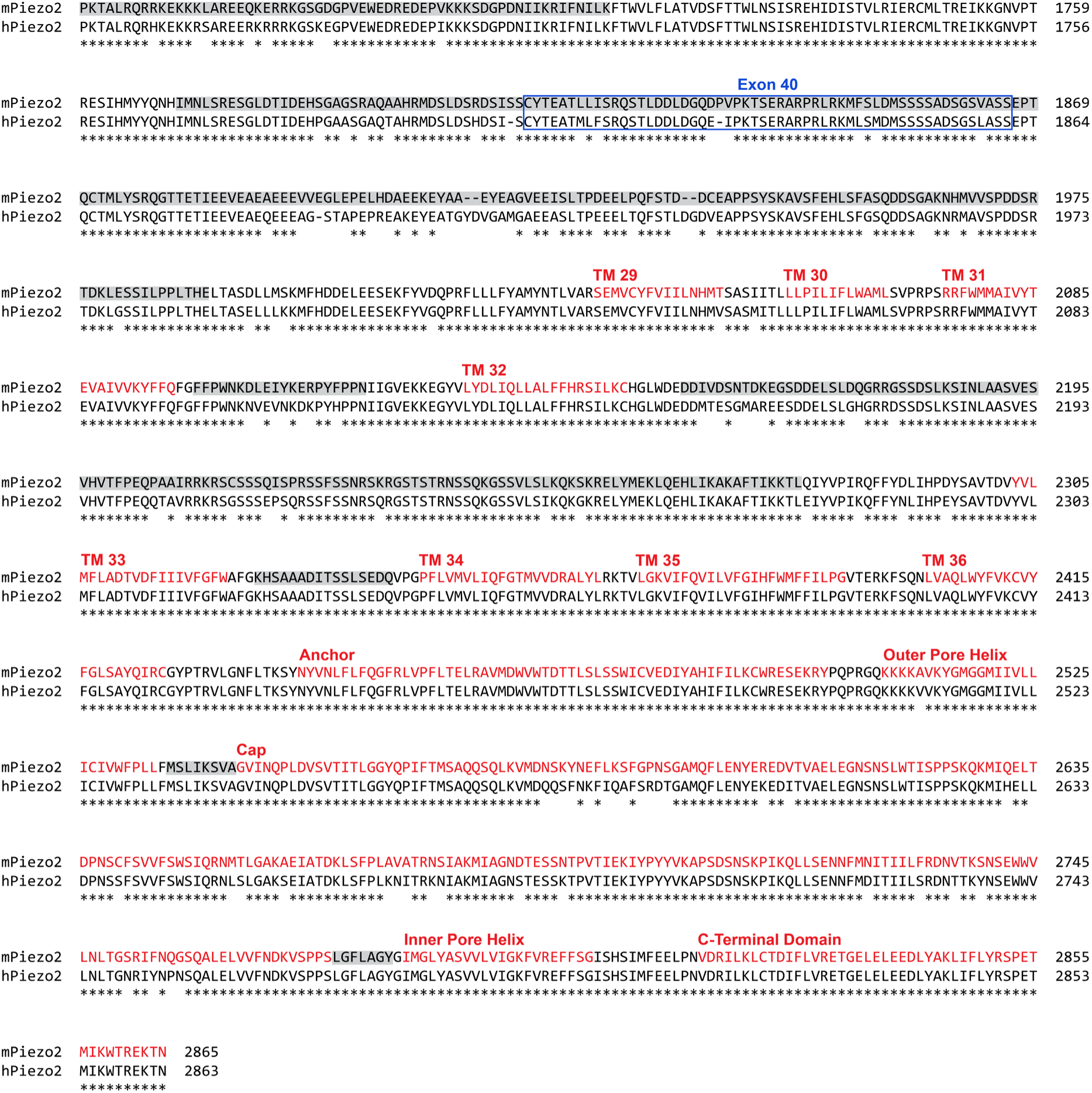
Sequence alignment and structural annotation of mouse and human Piezo2. Sequence alignment of mouse and human Piezo2 with all exons present. Alternatively spliced exons are highlighted by boxes, main structural features of mouse Piezo2 (PDB: 6KG7) are colored red, and domains that are not structurally resolved are shaded in grey.

**Supplemental Figure 2.**
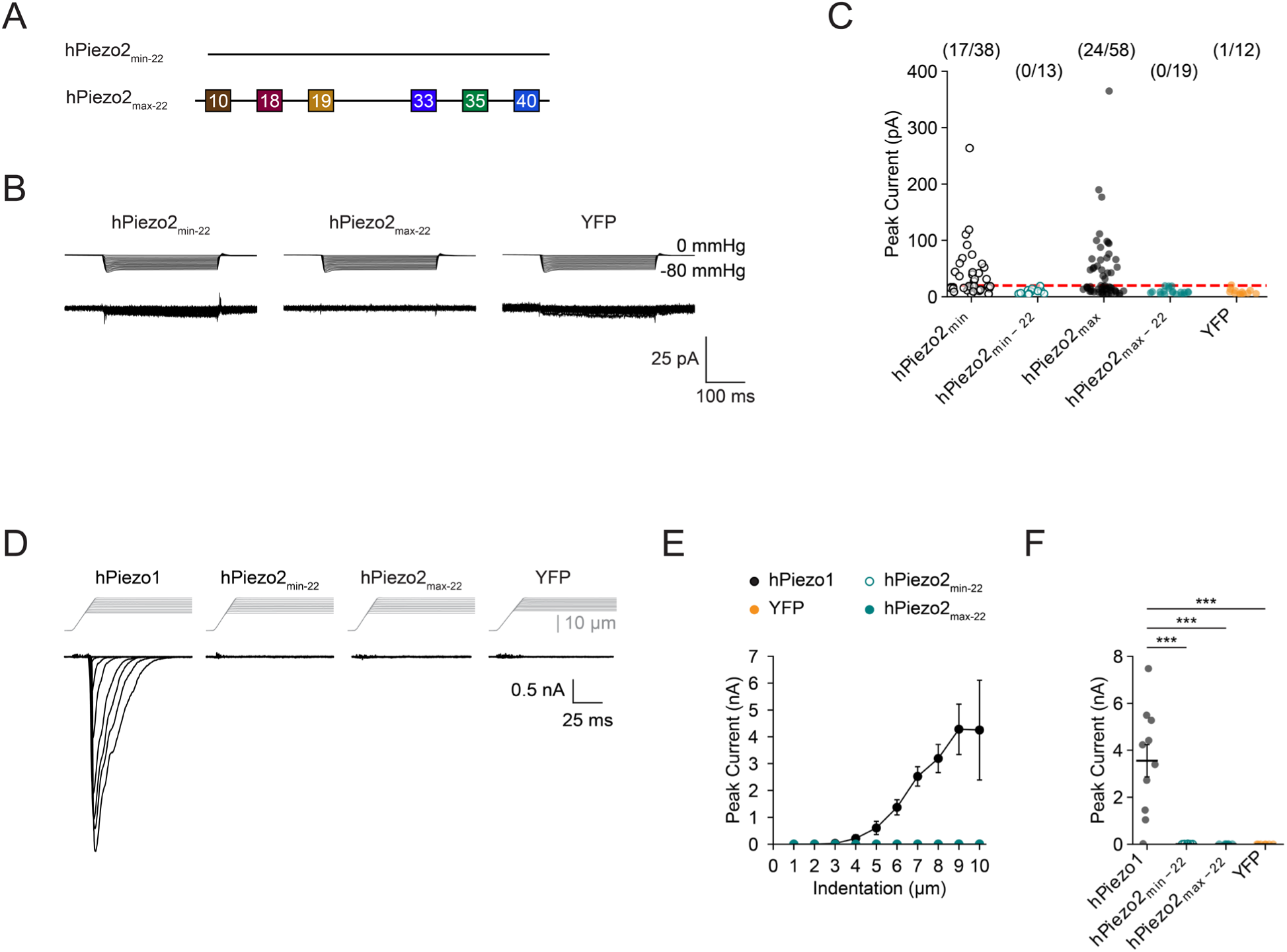
Removal of exon 22 from hPiezo2_max_ and hPiezo2_min_. ***A**, Schematic showing the alternatively spliced exons present in hPiezo2_min-22_ and hPiezo2_max-22_. **B**, Top: Pressure-step protocol from 0 to -80 mmHg (Δ = -5 mmHg) and representative currents recorded from Neuro2A-Piezo1ko cells heterologously expressing hPiezo2_min-22,_ hPiezo2_max-22_, or YFP. **C**, Peak current amplitudes obtained with the above protocol from cells expressing hPiezo2_min_, hPiezo2_min-22_, hPiezo2_max_, hPiezo2_max-22_, or YFP alone. The red dashed line illustrates the 20 pA threshold above which patches were used for analyzing tension responses. The number of patches with peak currents > 20 pA and the total number of patches are shown above. **D** Indentation-step protocol (Δ = 1 μm) and currents recorded from Neuro2A-Piezo1ko cells heterologously expressing hPiezo1, hPiezo2_min-22_, hPiezo2_max-22_, or YFP. **E**, Mean indentation response curves for hPiezo1 (n = 10), hPiezo2_min-22_ (n = 7), hPiezo2_max-22_ (n = 7), and YFP (n = 7). Data are represented as mean±S.E.M. Peak current amplitude values for all individual patches shown in (D). Bars indicate the mean±S.E.M. **F**, Mean values for peak current amplitude are as follows: hPiezo1 = 3,553±687 pA; hPiezo2_min-22_ = 10±1 pA; hPiezo2_max-22_ = 12±1 pA; YFP = 6±1 pA. Significance was determined using a one-way ANOVA (F = 16.2, p < 0.005) and Tukey’s HSD post-hoc comparison (hPiezo1/hPiezo2_min-22_, p < 0.005; hPiezo1/hPiezo2_max-22_, p < 0.005; hPiezo1/YFP, p < 0.005)*.

**Supplemental Figure 3.**
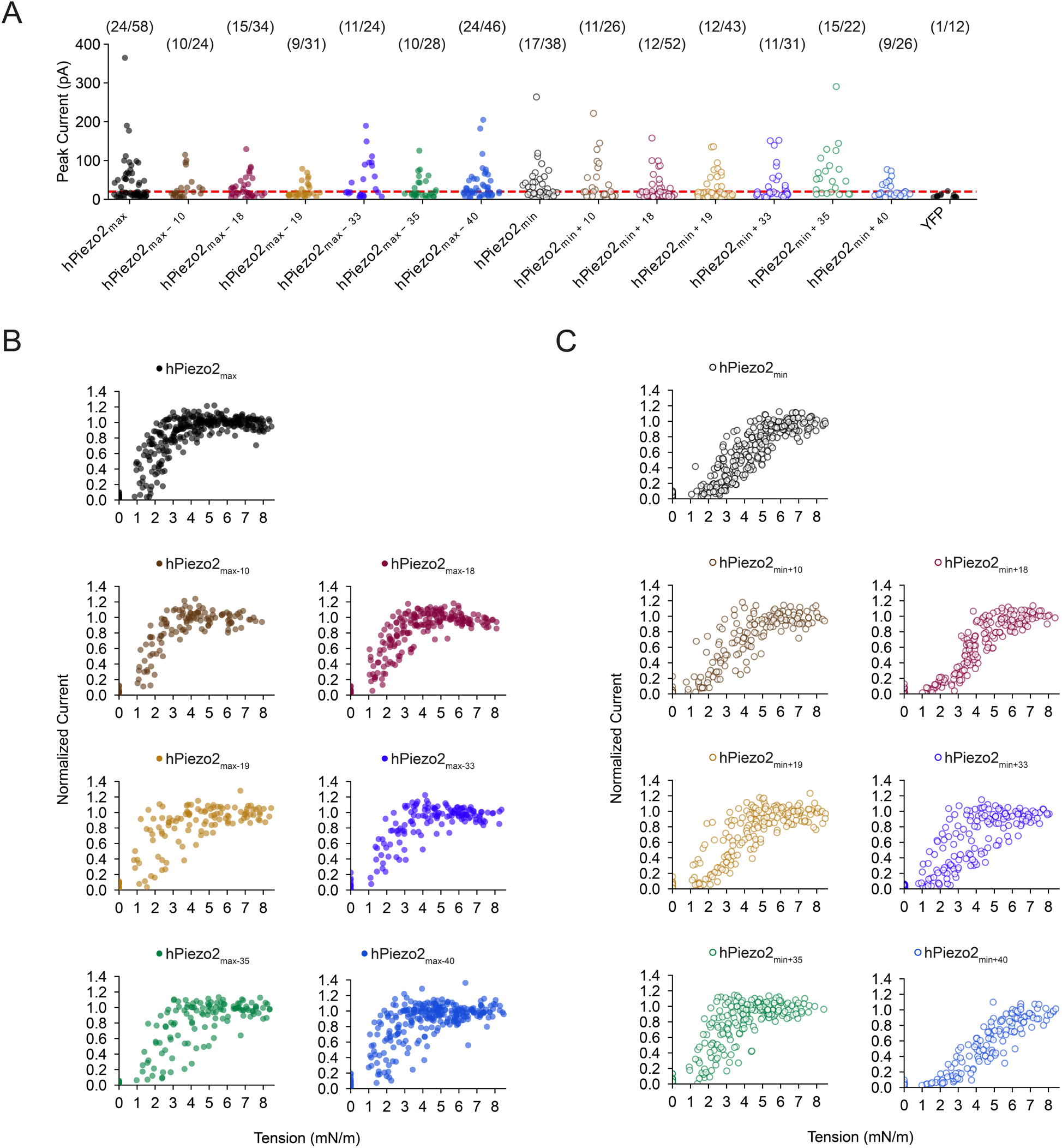
Peak currents and individual tension responses for hPiezo2_max(-)_ and hPiezo2_min(+)_ constructs. **A**, Peak current amplitudes obtained from Neuro2A-Piezo1ko cells heterologously expressing hPiezo2_max(-)_ and hPiezo2_min(+)_ constructs. The red dashed line illustrates the 20 pA threshold above which patches were used to calculate tension responses. The number of patches with a peak current > 20 pA and the total number of patches are shown above. **B**, Normalized tension response for all hPiezo2_max(-)_ constructs. Each point represents the tension and normalized current value elicited by one pressure step. **C**, Same as in (B), but for all hPiezo2_min(+)_ constructs.

**Supplemental Figure 4.**
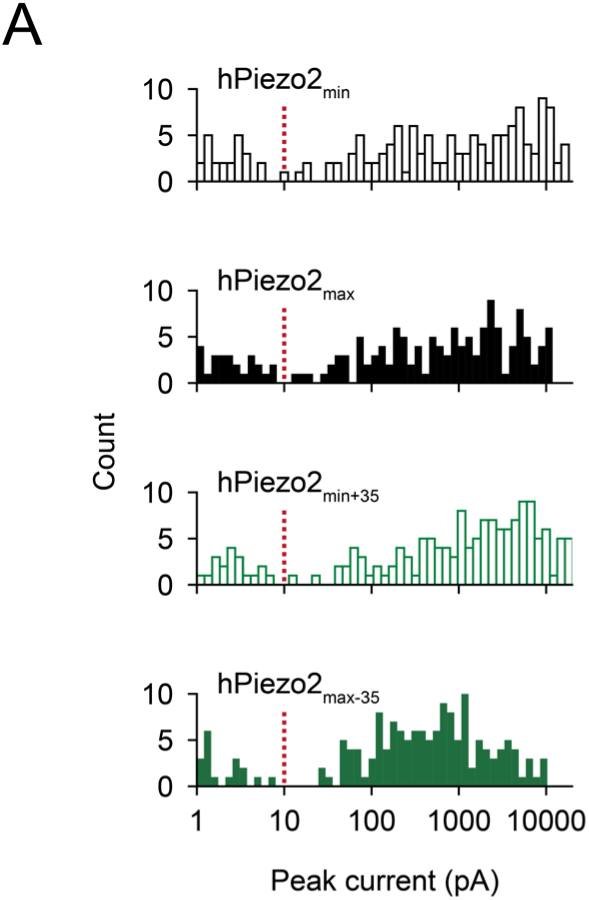
Indentation-induced peak current histograms for hPiezo2_min_, hPiezo2_max_, hPiezo2_min+35_, and hPiezo2_max-35_. ***A**, Histograms of peak currents from all indentation steps for each construct. The red dashed line illustrates the 10 pA peak current level, which was used as a response threshold*.

**Supplemental Figure 5.**
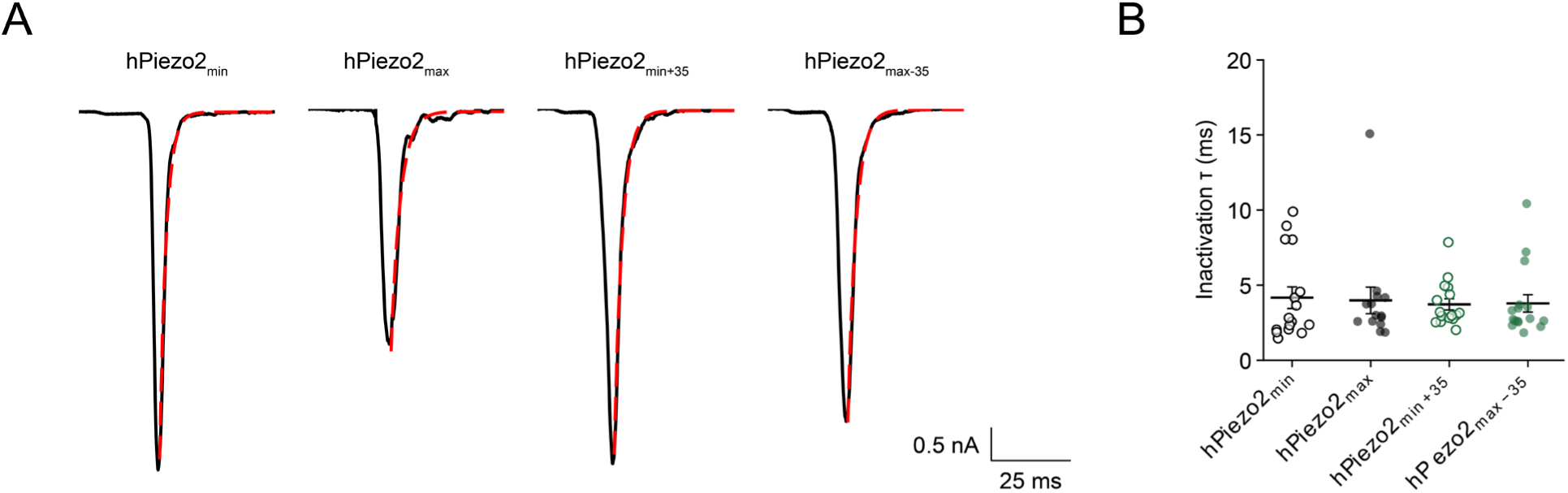
Inactivation kinetics for hPiezo2_min_, hPiezo2_max_, hPiezo2_min+35_, and hPiezo2_max-35_. **A**, Representative currents recorded from final indentation step for a whole-cell poke protocol from Neuro2A-Piezo1ko cells heterologously expressing hPiezo2_min_, hPiezo2_max_, hPiezo2_min+35_, or hPiezo2_max-35_. Currents were fit with a single exponential (red) and their rates of inactivation (τ) are follows: hPiezo2_min_ τ = 2.3±0.1 ms; hPiezo2_max_ τ = 3.0±0.1 ms; hPiezo2_min+35_ τ = 2.7±0.1 ms; hPiezo2_max-35_ τ = 2.6±0.1 ms. **B**, Inactivation time constants from single exponential fits to individual currents. Error bars indicate the mean±S.E.M. Mean values for inactivation (τ) are as follows: hPiezo2_min_ τ = 4.2±0.7 ms; hPiezo2_max_ τ = 4.0±0.9 ms; hPiezo2_min+35_ τ = 3.7±0.4 ms; hPiezo2_max-35_ τ = 3.8±0.6 ms. Significance was determined using a one-way ANOVA (F = 0.10, p = 0.96).

**Supplemental Figure 6.**
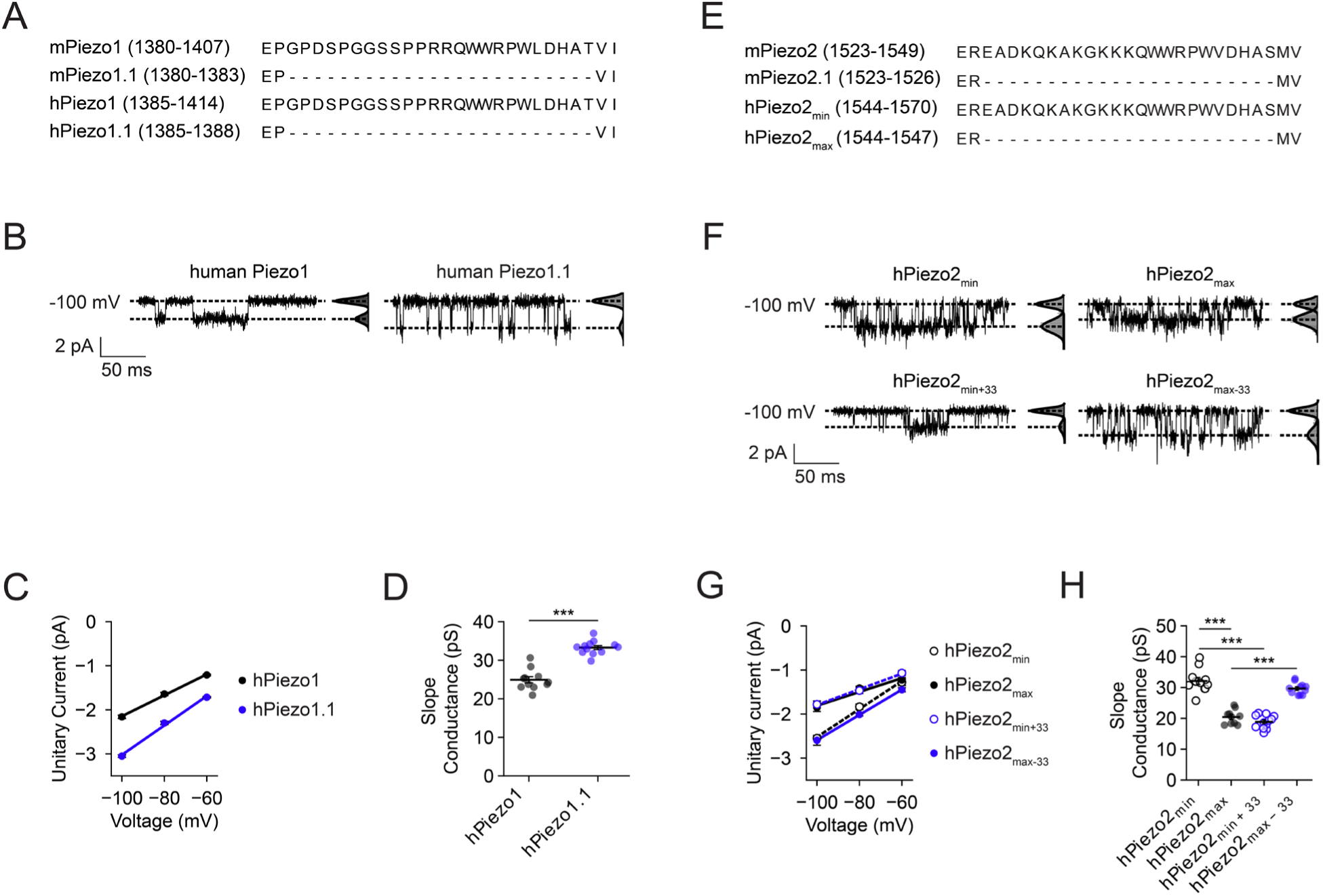
Single channel conductances of hPiezo1 and hPiezo2 constructs. **A**, Sequence alignment of mouse and human Piezo1 and Piezo1.1 that contain and lack exon 30, respectively. **B**, Representative cell-attached recordings from Neuro2A-Piezo1ko cells heterologously expressing hPiezo1 or hPiezo1.1 at a holding potential of -100mV and their current amplitude histograms. **C**, Representative current-voltage relationships generated from Neuro2A-Piezo1ko cells expressing hPiezo1 and hPiezo1.1. Slope conductances are g = 23.8±1.0 pS (hPiezo1) and g = 33.4±0.8 pS (hPiezo1.1). **D**, Slope conductances from all individual patches for hPiezo1 (n = 11) and hPiezo1.1 (n = 12). Error bars indicate mean±S.E.M. Mean slope conductances are as follows: hPiezo1 g = 24.9±0.8 pS; hPiezo1.1 g = 33.2±0.5 pS. Significance was determined using Welch’s unpaired t-test (t = 8.7, p < 0.0005). **E**, Sequence alignment of mouse Piezo2, mouse Piezo2.1., human Piezo2_min_, and human Piezo2_max_ that contain and lack exon 33, respectively. **F**, Representative cell-attached recordings from Neuro2A-Piezo1ko cells heterologously expressing hPiezo2_min_, hPiezo2_max_, Piezo2_min+33_, or Piezo2_max-33_, at a holding potential of -100mV and their current amplitude histograms. **G**, Representative current-voltage relationships generated from Neuro2A-Piezo1ko cells expressing hPiezo2_min_, hPiezo2_max_, Piezo2_min+33_, and Piezo2_max-33_. Slope conductances are as follows: hPiezo2_min_ g = 31.7±1.9 pS; hPiezo2_max_ g = 15.7±2.5 pS; Piezo2_min+33_ g = 17.8±2.3 pS; Piezo2_max-33_ g = 28.7±3.1 pS. **H**, Slope conductance values from all individual patches for hPiezo2_min_ (n =11), hPiezo2_max_ (n = 10), Piezo2_min+33_ (n = 12), and Piezo2_max-33_ (n = 11). Error bars indicate mean±S.E.M. Mean slope conductances are as follows: hPiezo2_min_ g = 32.1±1.2 pS; hPiezo2_max_ g = 20.4±0.7 pS; hPiezo2_min+33_ g = 18.9±0.6 pS; hPiezo2_max-33_ g = 29.7±0.6 pS. Significance was determined using a one-way ANOVA (F = 68.8, p < 0.0005) and Tukey’s HSD post-hoc comparison (hPiezo2_max_/hPiezo2_min_: p < 0.0005; hPiezo2_max_/hPiezo2_max-33_: p < 0.0005; hPiezo2_min_/hPiezo2_min+33_: p < 0.0005).

**Supplemental Figure 7.**
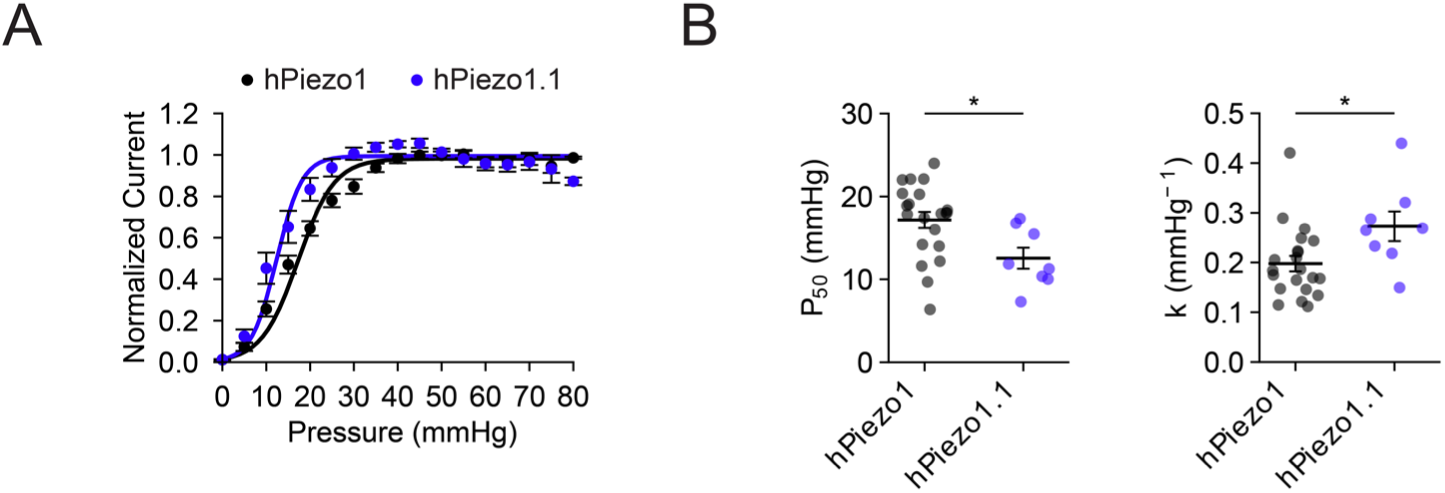
Pressure responses of hPiezo1 and hPiezo1.1. **A**, Average pressure response curves of hPiezo1 (n = 21) and hPiezo1.1 (n = 8). Data are plotted as normalized mean±S.E.M. **B**, Values of pressure of half-maximal activation (P_50_) and slope (k) from all individual patches. Error bars indicate mean±S.E.M. Mean values for P_50_ and k are as follows: hPiezo1 P_50_ = 17.2±1.0 mmHg, k = 0.20±0.02 mmHg^-1^; hPiezo1.1 P_50_ = 12.5±1.3 mmHg, k = 0.27±0.03 mmHg^-1^. Significance was determined using Welch’s unpaired t-test (P_50_ t = 2.9, p = 0.011; k t = 2.23, p = 0.048).

**Supplemental Figure 8.**
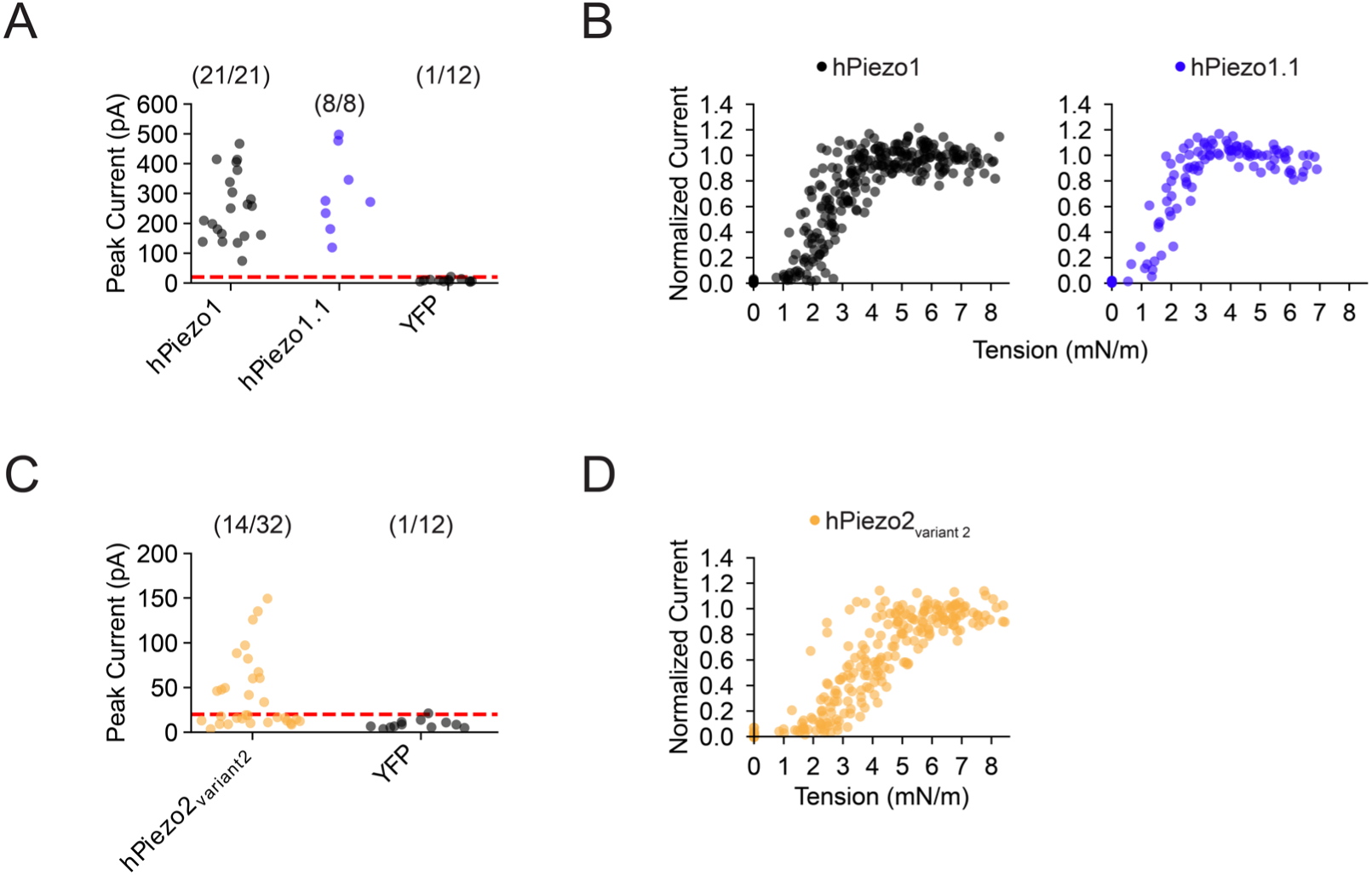
Peak currents and individual tension-responses of hPiezo1, hPiezo1.1, and hPiezo2 Variant 2. **A**, Peak current amplitudes from Neuro2A-Piezo1ko cells heterologously expressing hPiezo1 and hPiezo1.1. The red dashed line illustrates the 20 pA threshold above which patches were used for analyzing tension responses. The number of patches with a peak current > 20 pA and the total number of patches are shown above. **B**, Normalized tension responses. Each point represents the tension and normalized current value elicited by one pressure step. **C**, Same as in (A) but for hPiezo2 variant 2. **D**, Same as in (B) but for hPiezo2 variant 2.

**Table S1:**
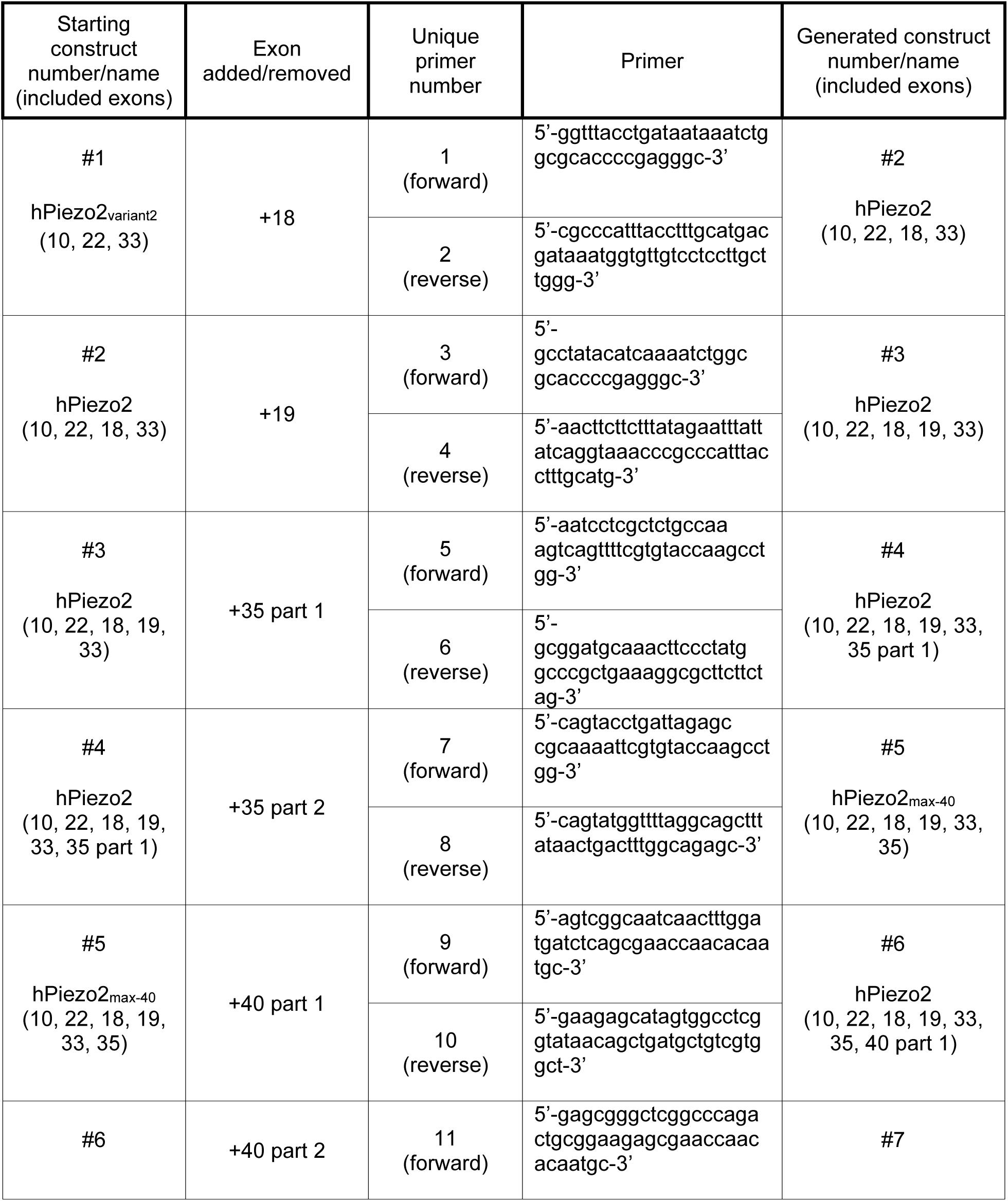
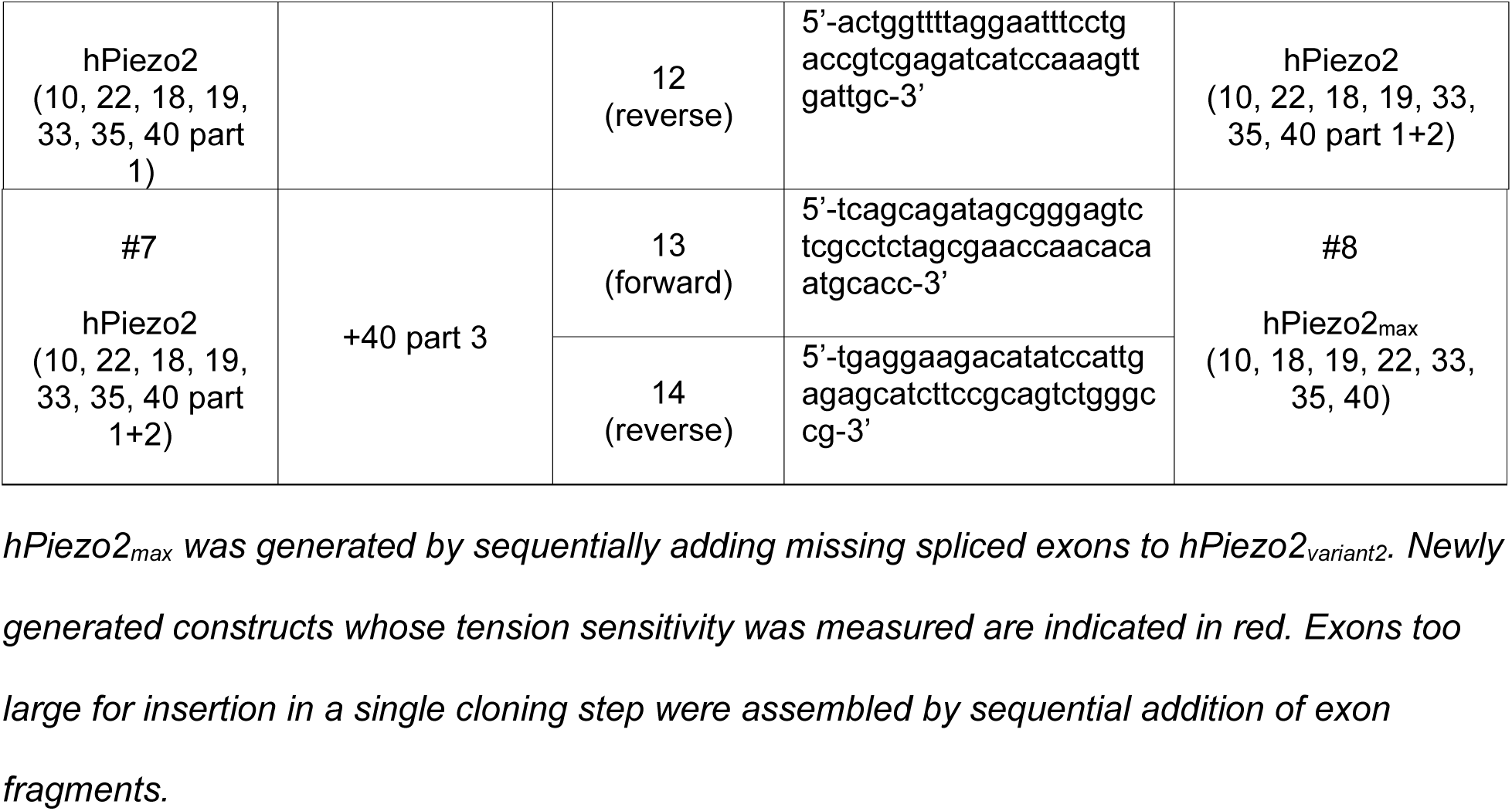
Primers for generating hPiezo2max starting from hPiezo2variant2 (hPIEZO2-pIRES2-mCherry-WPRE).

**Table S2:**
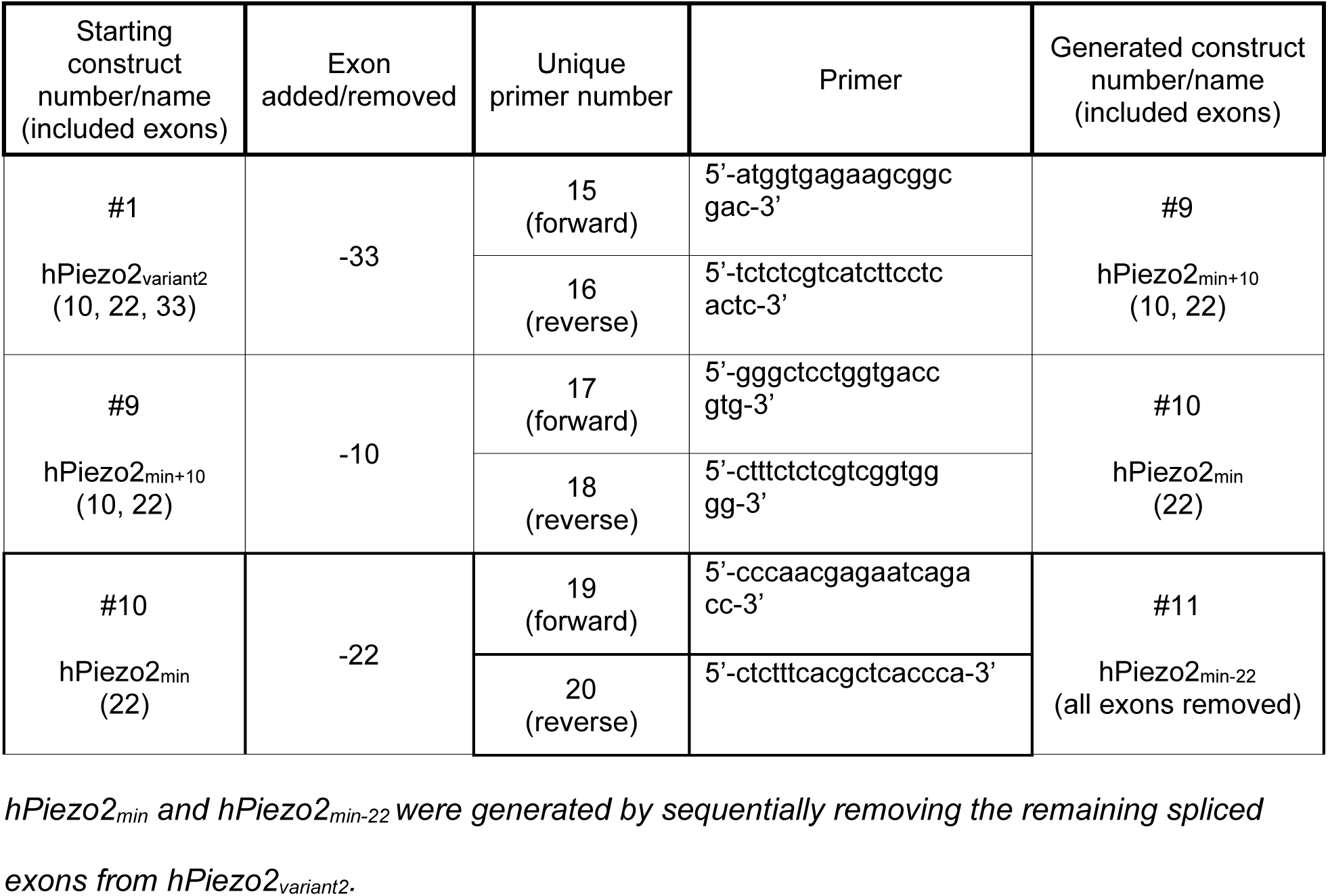
Primers for generating hPiezo2min and hPiezo2min-22, starting from hPiezo2variant2 (hPIEZO2-pIRES2-mCherry-WPRE).

**Table S3:**
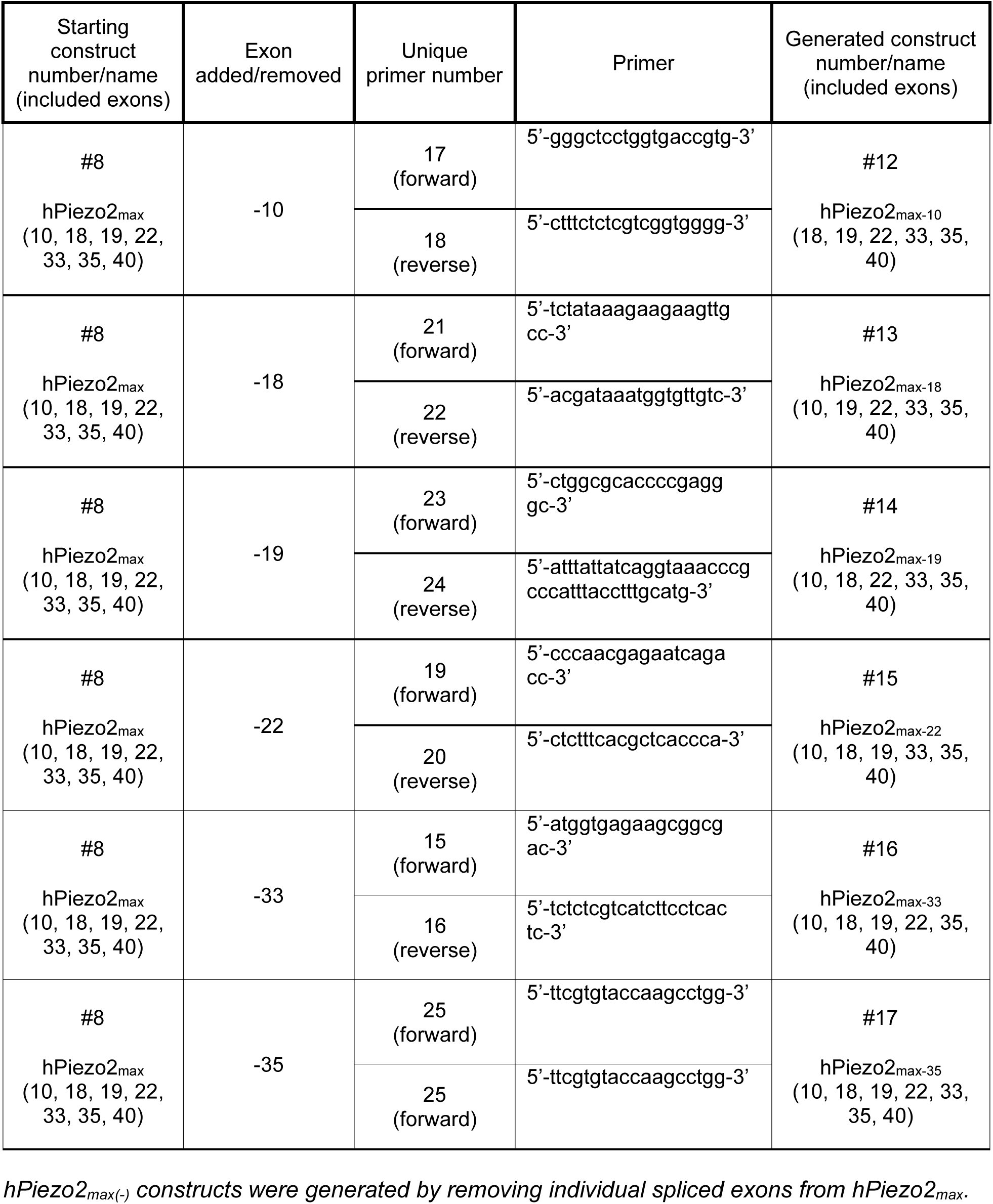
Primers for generating additional hPiezo2max(-) constructs, starting from hPiezo2max.

**Table S4:**
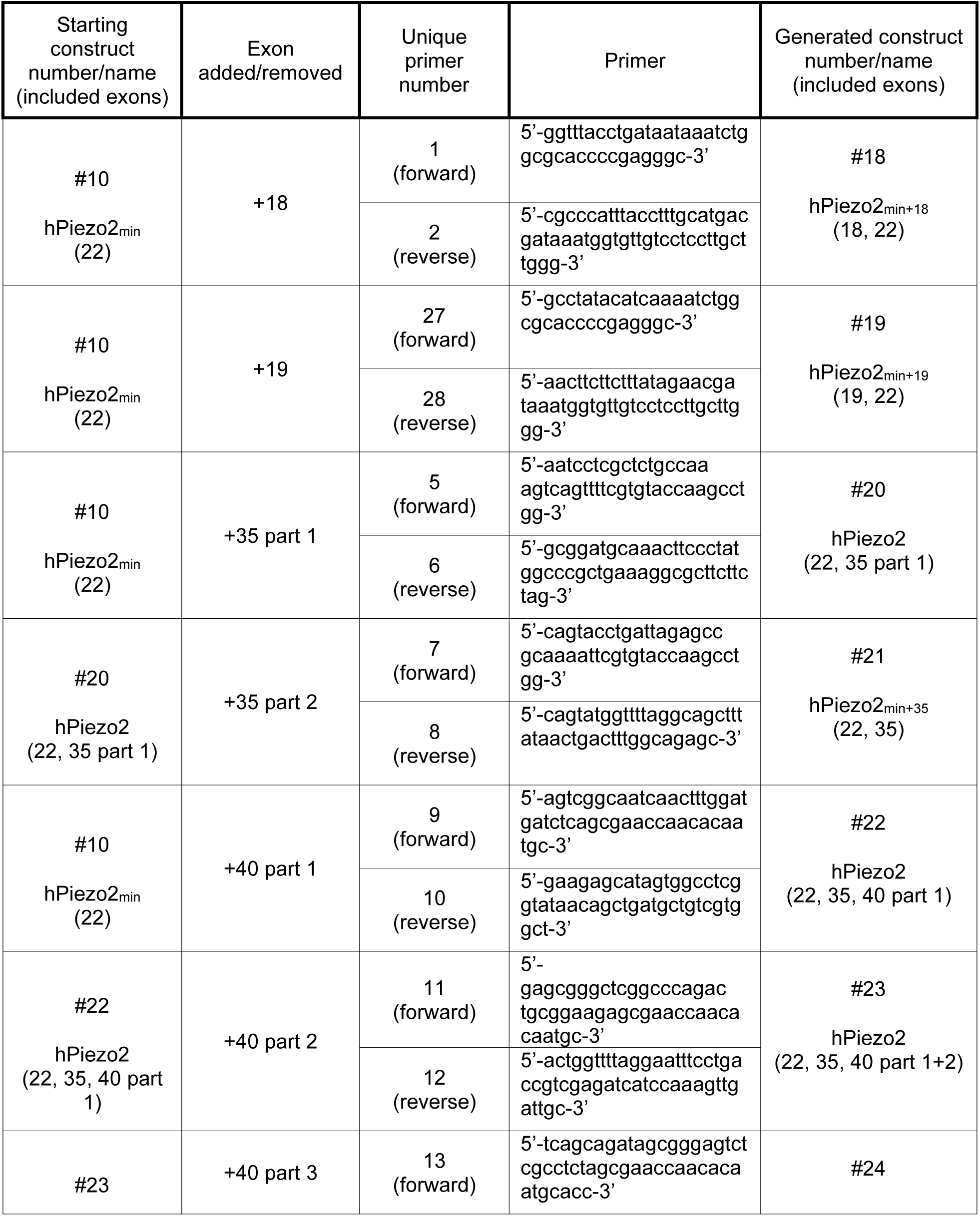
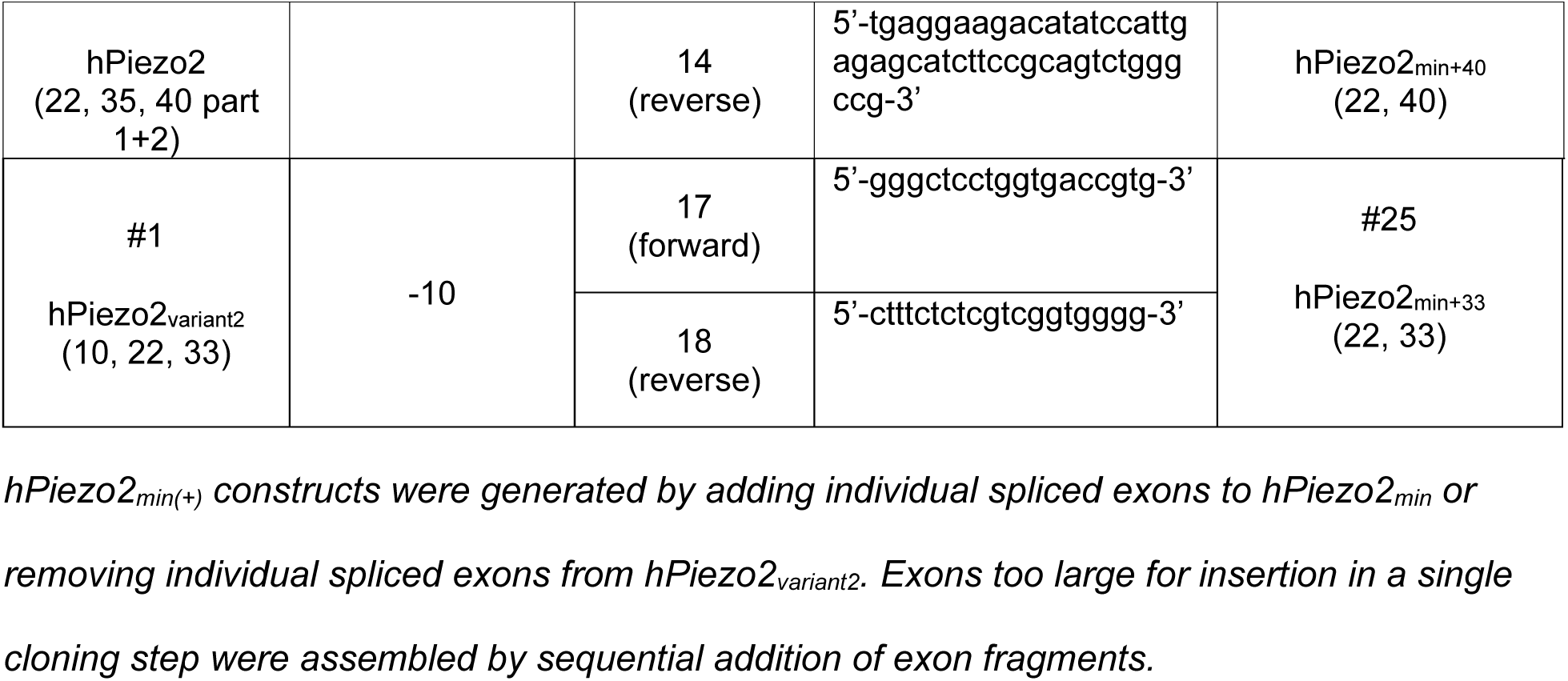
Primers for generating additional hPiezo2min(+) constructs, starting from hPiezo2min.

**Table S5:**
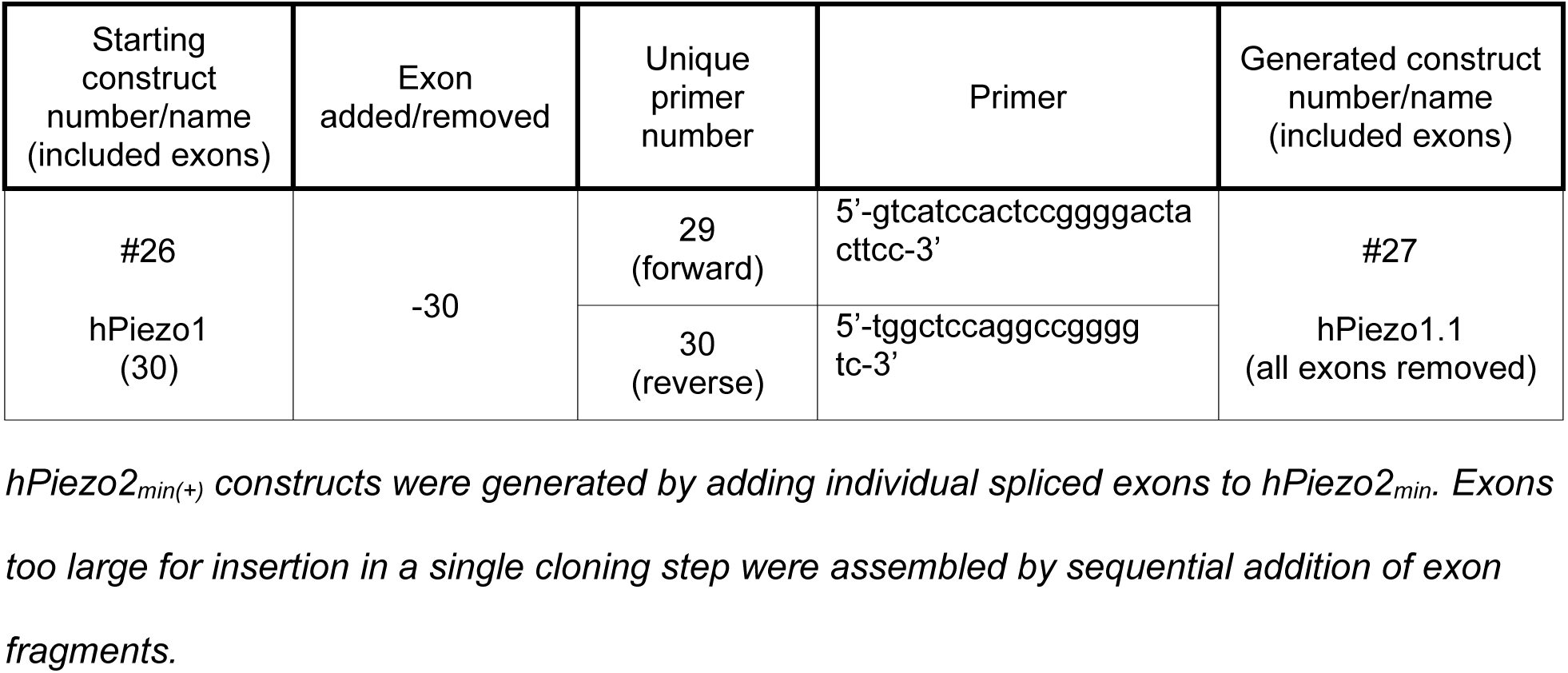
Primers for generating hPiezo1, starting from hPiezo1 (Human Piezo1-pIRES2-EGFP).

## Notes

https://github.com/GrandlLab

### Competing Interest Statement

The authors have declared no competing interest.

